# Reliability and disease sensitivity are dissociable properties of EEG foundation-model representations

**DOI:** 10.64898/2026.08.19.745052

**Authors:** Belay Tadesse Gebregergis, Haben Girmay Yhdego, Tewolde Teklu

**Author notes:** Belay Tadesse Gebregergis and Haben Girmay Yhdego contributed equally. **Corresponding author:** Belay Tadesse Gebregergis.

## Abstract

EEG foundation models (EEG-FMs) are evaluated almost entirely on disease-discrimination accuracy. A clinical biomarker additionally requires measurement reliability, the stability of repeated measurements on the same individual, which regulatory biomarker frameworks treat as a prerequisite that discrimination does not imply. We asked whether frozen EEG-FM representations provide such stability, whether it is predictable from conventional model descriptors, and what information supports it. We measured test-retest reliability, disease discrimination, and representation distinctiveness for nine frozen representations: six EEG-oriented foundation models spanning masked, contrastive and predictive pretraining, handcrafted spectral features, and two general-purpose time-series models with no EEG exposure. All were evaluated under one preprocessing pipeline across two healthy retest cohorts, at roughly one month and two years, and three neurodegenerative cohorts. Reliability was measured in healthy adults only; the disease cohorts contribute cross-sectional discrimination. Reliability, quantified as the intraclass correlation coefficient (ICC), varied enormously (mean 0.08 to 0.76; coeffiicient of variation, CV, 53.0%) while disease discrimination, measured as area under the receiver operating characteristic curve (AUC), occupied a far narrower range across the same nine (AUC CV 5.4%), a roughly tenfold difference in relative dispersion, described rather than formally tested. We did not test formal AUC equivalence, so we describe discrimination as varying substantially less than reliability rather than as equivalent. The variation was not consistently explained by pretraining paradigm or domain among the models studied, and a model with no EEG exposure was among the most reliable tested. Alpha-band information contributed disproportionately to reliability, whereas theta-band information ranked first for Alzheimer’s disease and frontotemporal dementia discrimination, directionally consistent with established EEG evidence in both conditions. Only the reliability half of that contrast is individually significant. Subspace geometry and band ablation, two methodologically distinct analyses, both indicate that the two properties are partially, not fully, dissociable, and network architecture determines whether the dissociation is preserved, traded off, or jointly degraded across depth. Reliability showed no detectable association with discrimination, pretraining paradigm, or domain, and had to be measured directly. We recommend it become a standard evaluation axis for EEG-FM representations intended for longitudinal or biomarker use, and release a reproducible pipeline.

## 1. Introduction

A model can distinguish patients from controls without providing a stable measurement of the individual. EEG foundation models, large self-supervised networks pretrained on heterogeneous EEG such as LaBraM (Jiang et al., 2024) and EEGPT (Wang, G. et al., 2024), are increasingly proposed as the representational substrate for clinical EEG biomarkers. Their evaluation, however, has centred almost entirely on the former property: classification accuracy and transfer performance. Recent critical syntheses argue this evaluation is heterogeneous and limited, leaving real-world utility unclear (Kuruppu et al., 2026; Xiong et al., 2025). Comprehensive benchmarks add that linear probing of frozen embeddings is often insuffiicient, that specialist models remain competitive, and that the relationship between scale and downstream generalisation is not yet consistent across benchmarks (Liu et al., 2026; Yang et al., 2026).

This pattern recurs across biomarker modalities. Across neuroimaging, from MRI morphometry to fMRI connectivity “fingerprinting” (Finn et al., 2015), between-group separability is typically established long before within-individual stability, even though stability is what governs whether a marker can track change over time. A decade of fMRI test-retest work makes the point quantitatively: individual connectivity measurements have a mean ICC near 0.29 (Noble et al., 2019), and a meta-analysis of 90 task-fMRI experiments found a mean ICC of 0.397, low enough that the authors concluded such measures are not currently suitable for biomarker discovery (Elliott et al., 2020). Those numbers show both that reliability is measurable and that representations widely used for group-level inference can be only modestly reliable in the individual.

There is a deeper reason to expect the two properties to come apart, and it is not specific to neuroimaging. Hedge, Powell and Sumner (2018) named it the reliability paradox: the experimental effects that replicate most robustly at the group level are precisely those with small between-subject variance, and small between-subject variance is what destroys reliability for individual differences. An effect can be highly reproducible as a group contrast and nearly useless as an individual measurement, for a reason internal to how the two quantities are defined. The same tension appears in prognostic modelling, where Riley and Collins (2023) show that a clinical prediction model’s estimated risks can be considerably unstable at the individual level even when the model performs acceptably in aggregate, and argue that stability should be examined at the individual level rather than inferred from group-level performance. That our results take the same shape in learned EEG representations suggests the problem belongs to predictive modelling generally rather than to EEG foundation models specifically. Discrimination rewards between-group variance. Reliability requires within-individual stability relative to between-individual spread. A monitoring biomarker, in the FDA-NIH BEST terminology, is one measured repeatedly to assess the status of a disease over time (FDA-NIH Biomarker Working Group, 2016), which means it must return consistent values on repeated measurement of the same person. We therefore treat stability, rather than group separability, as the property that governs longitudinal use. Recent reviews have highlighted the narrow scope of current EEG-FM evaluation and called for more practically relevant evaluation axes (Kuruppu et al., 2026); test-retest reliability is one such axis, and it is the one we operationalise here. Yet to our knowledge, no prior EEG-FM evaluation has foregrounded embedding test-retest reliability, asked whether it differs across architectures, or asked whether the same representational information supports both properties. This omission does not reflect a shortage of reliability data in EEG itself. Test-retest reliability of resting-state EEG has been quantified across spectral power, microstates, and source-space connectivity in both young and older adults (Popov et al., 2023). The spectral-parameterisation features we adopt in our handcrafted baseline, the aperiodic exponent and offset, reach ICC values above 0.70 across sessions spaced 90 minutes and 30 days apart in healthy young adults (McKeown et al., 2024). Individual alpha peak frequency is stable enough to be treated as a trait marker, unchanged across roughly 100 hours of cognitive training in both younger and older adults (Grandy et al., 2013). Over an interval comparable to our own longer retest cohort, posterior alpha relative power reaches an ICC of 0.843 and alpha peak frequency an ICC of 0.734 across five years in healthy adults (Park et al., 2026). These values give external reference points against which the handcrafted-feature reliability we report below can be judged. What has not been done is to turn that same measurement lens on learned representations.

The distinctiveness axis has its own precedent. Individuals can be identified from resting EEG, most accurately from the aperiodic component of the power spectrum rather than from canonical band power (Demuru & Fraschini, 2020). Brief segments of resting neurophysiological activity are enough for individual differentiation in MEG (da Silva Castanheira et al., 2021), the electrophysiological counterpart of connectome fingerprinting (Finn et al., 2015). Foundation models are best read as the current stage of a longer line of self-supervised EEG representation learning (Banville et al., 2021), not a break from it.

A cluster of concurrent audits sharpens the motivation for measuring reliability and distinctiveness separately, and each converges on the same underlying worry from a different direction. Lin et al. (2026) show that subject-identity information can substantially influence frozen EEG-FM representations and confound clinical decoding. Zare (2026a) finds that frozen-probe benchmarking with targeted negative controls makes classical features competitive with, or better than, pretrained embeddings on several clinical tasks, and that dataset identity is decodable from frozen embeddings at near-ceiling accuracy. In a companion analysis, Zare (2026b) reports a related spectral-temporal dissociation in five EEG-FMs, discussed in Section 4. Širca et al. (2026) evaluate robustness, interpretability, and expressiveness rather than accuracy alone, and find that much of the apparent weakness of frozen EEG-FM representations is attributable to pooling rather than to the representations themselves, a point we return to when interpreting our own layer-wise results. Tang et al. (2026) probe which handcrafted features are recoverable from EEG-FM representations layer by layer, and report that the depths at which those features are most strongly encoded cluster in the lower-to-middle backbone rather than at the final layer. Large standardised benchmarks have also begun to include general-purpose time-series models alongside EEG-specific ones, and report that EEG-specific models do not consistently outperform them (Kontras et al., 2026). What these audits share is the finding that accuracy on a clinical label can coexist with representational properties that would disqualify the same embedding from clinical use. Several of the individual techniques we use are established. Frequency-band ablation of frozen embeddings has been used to localise the spectral basis of disease discrimination (Zare, 2026a), layer-wise probing has been used to show that the final layer is not where task-relevant information is richest (Širca et al., 2026; Tang et al., 2026), and general-purpose time-series models have been benchmarked on EEG classification (Kontras et al., 2026). What has not been done, to our knowledge, is to apply any of these to test-retest reliability of frozen representations, measured as the stability of the same quantity on repeated recording of the same person. We distinguish this from the reliability-named task families in recent benchmarks (Lu et al., 2026), which score cross-session subject identification rather than measurement stability. Our contribution is not to introduce reliability or identifiability as concepts, but to measure them directly and jointly, under a single preprocessing pipeline, across a representative set of frozen EEG foundation models, and then to ask what information supports each. The comparison that follows from doing both at once, between the information that supports reliability and the information that supports discrimination, is the part we believe is new.

We evaluate EEG representations on three partially separable axes: discrimination, or between-group separability; measurement stability, or within-individual consistency on retest as quantified by ICC; and representation distinctiveness, whether an individual is identifiable from their representation, which can dissociate from stability. We study six frozen foundation models spanning the dominant self-supervised paradigms: masked or tokenized reconstruction (LaBraM, CBraMod (Wang, J. et al., 2025), and BIOT (Yang, C. et al., 2023)), contrastive alignment (EEGPT and BENDR (Kostas et al., 2021)), and joint-embedding prediction (SignalJEPA; Guetschel et al., 2024). Together with handcrafted spectral features, these give seven EEG-oriented representations in total. We add two general-purpose time-series foundation models with no EEG exposure, MOMENT (Goswami et al., 2024) and Chronos (Ansari et al., 2024), to ask a complementary question: is EEG-specific pretraining necessary for reliability at all?

This paper asks four questions in sequence, each motivating the next. First, does reliability vary, and is that variation predictable from discrimination performance, pretraining paradigm, or pretraining domain? Second, is the variation real, structurally present in the trained representations, rather than an artifact of dimensionality or acquisition conditions? Third, what information supports reliability, and is that information inherited trivially from whichever raw-signal feature happens to be most reliable, or is there a more specific explanation? Fourth, is the information that supports reliability the same information that supports disease discrimination, and if not, where in a model’s architecture does this distinction live?

We combine two classes of evidence throughout. Correlational and geometric analyses, including variance decomposition, representation drift, linear centred kernel alignment (CKA; Kornblith et al., 2019) and singular vector canonical correlation analysis (SVCCA; Raghu et al., 2017), dimension-physiology correlation, subspace-overlap geometry between the subject-stable and disease-discriminative directions of the embedding space, and a supervised-contrastive recoverability probe, use existing frozen embeddings and require no new model inference. Perturbational analyses manipulate the input signal, readout layer, or recording quality and re-run inference: frequency-band ablation via zero-phase band-stop filtering before re-embedding, layer-wise reliability via forward hooks on intermediate transformer or convolutional blocks, AUC-underband-ablation on disease cohorts, and acquisition-degradation stress tests. These provide evidence one level closer to causal than correlation alone. One caveat applies throughout: band-stop filtering is perturbational, not causal in the interventionist sense, for the reasons set out in Section 5.

## 2. Methods

This section describes the data, the preprocessing applied to it, the nine representations compared, and the statistical procedures used to quantify reliability, discrimination, and their relationship. The design principle throughout is that every representation sees the identical signal. One preprocessing pipeline produces one clean continuous recording per session, and that recording is the sole input to all nine representations and to every perturbational analysis, so that differences between representations cannot be attributed to differences in how their inputs were prepared.

### 2.1 Datasets

All data are pre-existing, de-identified, publicly available EEG recordings. No new human-subjects data were collected. Five public cohorts were used; acquisition details are in **Supplementary Table S1**. **Wang** (OpenNeuro ds004148; Wang et al., 2022) contributed 60 healthy adults across three eyes-closed resting sessions, two within about 90 minutes of each other and one about a month later, recorded at 61 channels and 500 Hz. This is the short-interval healthy retest anchor. **Henao Isaza** (ds007176; Henao Isaza et al., 2026a) contributed 45 healthy adults across up to four eyes-closed sessions spanning about two years, recorded at 60 channels and 1000 Hz; 39 of the 45 had two or more usable sessions. The cohort’s own handcrafted-feature test-retest reliability, computed under a different automated preprocessing pipeline, is reported by Henao Isaza et al. (2026b), which provides an external reference point for the handcrafted end of our range in this cohort. This is the long-interval healthy retest anchor. **Miltiadous** (ds004504; Miltiadous et al., 2023) is published as 36 AD, 23 FTD, and 29 HC recordings. After this study’s unified quality-control pipeline (Methods 2.2), 34 AD, 22 FTD, and 29 HC recordings had usable exports for the discrimination analyses, giving 85 recordings in total and 63 for the AD-versus-HC contrast specifically, recorded at 19 channels. **Rockhill** (ds002778; Rockhill et al., 2021) and **Cavanagh** (ds003490; Cavanagh, 2021; Cavanagh et al., 2018) are two independent Parkinson’s disease cohorts, each contrasting patients and controls recorded off medication. Control age distributions matched the disease groups in every cohort, with all mean-age differences under five years; age and sex were included as additional predictors rather than partialled out, so they contribute to the reported AUCs equally across representations, and age and sex alone reach an AUC of about 0.64 in the Miltiadous contrasts and below chance in both Parkinson’s cohorts (**Supplementary Table S18**); and MMSE scores confirmed the expected group separation (AD 17.7, FTD 22.1, controls 30.0, PD cohorts approximately 29). All analyses used frozen embeddings, with no fine-tuning of any model.

### 2.2 Unified preprocessing

One pipeline applied identical quality control to every representation. A recording excluded by these rules was excluded from every representation, not just some. Standard 10-20 and 10-10 channels were retained. Bad channels were detected using a scale-invariant amplitude criterion and interpolated by spherical spline, and a recording was excluded if more than 30% of its channels were bad. Each recording was unit-normalised, band-pass filtered from 1 to 45 Hz, and re-referenced to the average. Data were epoched into 4-second windows with 150 microvolt peak-to-peak rejection, and a recording was retained only if at least 50% of its epochs survived and at least 15 seconds of clean data remained. The result was a clean continuous signal that fed every representation, including the perturbational re-embedding analyses described below. Those analyses reuse this exact clean signal and the exact embed_*_from_clean() functions from the same pipeline, so any perturbational effect can be attributed to the input manipulation itself rather than to a different embedding procedure.

### 2.3 Representations

Nine representations were derived from the identical clean signal. Handcrafted features (13 total) covered absolute and relative band power from delta through gamma, the aperiodic exponent and offset, and individual alpha peak frequency. Six foundation models were used, all frozen and loaded via braindecode (Schirrmeister et al., 2017): LaBraM-Base (200 dimensions), CBraMod (200 dimensions), and BIOT (256 dimensions) for masked or tokenized reconstruction; EEGPT (2048 dimensions) and BENDR (512 dimensions) for contrastive alignment; and SignalJEPA (64 dimensions) for joint-embedding prediction. Each model has its own window and montage conventions, detailed in **Supplementary Table S2**. Together with handcrafted features, this gives seven EEG-oriented representations; the remaining two are the general-purpose time-series models described below, for nine representations in total.

For BENDR and SignalJEPA, the distributed downstream projection head produced near-constant output across recordings under frozen inference, with mean pairwise cosine distance close to zero, so neither head could distinguish one individual from another. We verified this before computing any reliability metric (**Supplementary Tables S2 and S3**) and used each model’s convolutional feature-encoder representation instead, global-average-pooled over time. This was a principled, uniformly applied choice, made and validated before any downstream result was computed, not selected afterward to produce one.

Two general-purpose time-series foundation models with no EEG exposure, MOMENT-1-large (1024 dimensions, masked reconstruction; Goswami et al., 2024) and Chronos-T5-small (Ansari et al., 2024) (512 dimensions, an encoder-decoder architecture using scaling-and-quantisation to-kenisation), were embedded identically to test whether EEG-specific pretraining is necessary for reliability at all. Each channel was treated as an independent univariate series, resampled to 200 Hz, z-scored, windowed, embedded, and mean-pooled.

### 2.4 Reliability, discrimination, and their relationship

#### Reliability

Test-retest reliability was quantified as ICC(2,1), the two-way random-effects, absolute-agreement form for a single measurement, computed per handcrafted feature and per embedding dimension using each subject’s longest available retest interval. For a single measure, ICC(2,1) equals 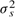 divided by the sum 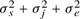 (Shrout & Fleiss, 1979), where 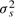 is between-subject variance, 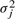 is between-session variance, and 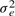 is residual error. Following the reporting guidance of Koo and Li (2016), we state the form explicitly because the ten ICC variants are not interchangeable and are frequently reported without specification. We chose this absolute-agreement form over a consistency-only form such as ICC(3,1) because a representation intended to support longitudinal biomarker tracking must reproduce comparable absolute values on retest, not merely preserve the relative ordering of subjects. A systematic session-to-session shift, which a consistency-based ICC would tolerate, would still constitute unreliability for this purpose. Sessions were treated as a random rather than fixed factor because the retest occasions in each cohort are a sample of possible remeasurement times, not a fixed set of conditions of intrinsic interest. This choice of metric makes the paper’s central dissociation explicit even at the level of the formula: discrimination rewards large between-subject variance alone, but reliability additionally penalises between-session and residual variance, so a representation can discriminate well while still being unreliable if its session and error variance are also large.

Dimensionality-agnostic multivariate metrics were computed alongside the per-dimension mean ICC, so that no ordering claim rests on averaging over correlated, non-identifiable or arbitrarily scaled latent dimensions. For embeddings *E_i_* with subject labels *s_i_*, the ratio R is the mean distance between same-subject pairs divided by the mean distance between different-subject pairs, over all recording pairs in a cohort; lower values indicate that an individual’s sessions cluster tightly relative to other people’s. R is reported under two distances, because they answer different questions. The consistency ratio uses correlation distance, *d*(*E_i_*, *E_j_*) = 1 minus the correlation between *E_i_* and *E_j_*, which is invariant to shifting or rescaling a whole recording and therefore asks only whether the pattern across dimensions is reproduced. The agreement ratio uses Euclidean distance on cohort-standardised dimensions, which a session-to-session offset does move. Only the second is the multivariate counterpart of the absolute-agreement ICC(2,1) used throughout this paper, since ICC(2,1) also penalises systematic session shifts; the first is a consistency-type measure and is reported for continuity with the earlier version of this analysis. Where the two disagree, the agreement ratio is the one comparable to the headline ICC. Ordering claims are corroborated by convergence between the ICC and the agreement ratio in both cohorts, and further by a matched-dimensionality PCA control that does not depend on either choice (Section 3.1).

Representations were compared using a paired subject-level bootstrap: 2000 iterations for the original benchmark comparisons, and 1000 iterations for the mechanistic follow-up’s band-ablation and layer-wise analyses, which involve many more model-by-condition combinations and were scaled down for tractability. Both used a fixed seed of 42. Every pairwise representation comparison, with its confidence interval, is tabulated in **Supplementary Table S4**. Results are reported as 95% confidence intervals against a pre-specified smallest-effect-size-of-interest of 0.05, rather than as binary significance decisions. This limits, though does not eliminate, the multiplicity concern that comes from testing many representation pairs. No single pairwise comparison carries this paper’s central claims; those instead rest on convergence across the full reliability ordering, the dimensionality-agnostic metric, and the robustness analysis described in Section 3.1.

#### Discrimination

For each disease-versus-control contrast, we fit an L2-regularised logistic regression with repeated, ten-times stratified 5-fold cross-validation, including age and sex as covariates where available. Significance was assessed against a permutation-shuffled null, using 200 permutations by default and fewer for specific high-dimensional or compute-constrained passes; these reductions are flagged inline and recorded per row in each results file’s n_perm_used column.

#### Statistical convention for multiple comparisons

A single correction across every test in the study would treat structurally different comparisons, planned at different times for different purposes, as one family, while no correction at all would understate the multiplicity risk. We took a middle path: we defined comparison families based on how the tests were structured within the manuscript, not registered in advance of data collection, since this study is exploratory and not pre-registered (Limitations), and applied Benjamini-Hochberg false-discovery-rate correction at q < 0.05 within each family. Per-test values for every family are in the released tier2_multiple_comparisons_bh_fdr.csv. Family 1 covers the original 36 representation-by-contrast discrimination tests (**Supplementary Table S5**). FDR correction left the set of significant results unchanged, at 27 of 36 significant both before and after correction. Bonferroni correction, at *α* = 0.05/36 ≈ 0.0014, is not informative here because it falls below this permutation test’s resolution floor of roughly 1/201 ≈ 0.005 with 200 permutations, so we report the FDR result instead. Family 2 covers the AD-versus-HC band-ablation AUC permutation tests, 18 tests spanning baseline, no-alpha, and no-theta conditions across six representations. All 18 remain significant after correction, at a BH cutoff of p ≤ 0.0199. Family 3 covers the FTD-versus-HC band-ablation AUC permutation tests, 36 tests spanning all five bands across six representations. Twenty-nine of 36 remain significant after correction, at a BH cutoff of p ≤ 0.0398. The seven that lose significance are BENDR’s no-alpha, no-beta, no-gamma, and no-theta conditions, CBraMod’s no-beta and no-theta conditions, and LaBraM’s no-theta condition. These losses concentrate in BENDR, whose band-ablation reliability effects are also uniformly null (Section 3.2), and do not touch the baseline or no-theta comparisons for the other five representations that carry the headline theta-driven-discrimination claim in Section 3.3.

Families 4 and 5 cover the layer-wise comparisons. Because the layers being compared are measured on the same subjects, both use a paired subject-level bootstrap: within each draw one resampled subject index is applied to both layers and the statistic is their difference. Two-sided p-values are twice the smaller tail proportion with the conventional plus-one correction, giving a resolution floor of 0.002 at 1000 draws and 0.004 at 500. Family 4 is the three layer-wise reliability contrasts we make claims about, BENDR’s first against final encoder stage in each cohort and LaBraM’s block 8 against its final block, at 1000 draws. Family 5 is the three pairwise LaBraM layer-wise AUC contrasts, at 500 draws. Benjamini-Hochberg correction is applied within each family, and all draws are released so both can be recomputed.

The “roughly tenfold” comparison in the Abstract and Section 3.1 is the ratio of coeffiicients of variation, computed over the same nine representations for both quantities. Reliability is Wang mean ICC (mean 0.455, population SD 0.241, CV 53.0%). Discrimination is each representation’s mean AUC across the three adequately powered contrasts, AD-versus-HC, FTD-versus-HC, and Cavanagh-PD (mean 0.789, population SD 0.042, CV 5.36%). The ratio is 53.00 divided by 5.36, or 9.9.

This ratio is not sensitive to the choice of contrasts. SignalJEPA’s failure on the PD cohorts (Section 3.3) is the single largest contributor to discrimination spread, so restricting the AUC average to the AD and FTD contrasts, where all nine representations discriminate significantly, narrows discrimination further (mean 0.814, CV 3.8%) and widens the ratio to 13.8. We report the three-contrast figure because it is the more conservative of the two and because it matches the axis plotted in **Figure 1**.

**Figure 1.**
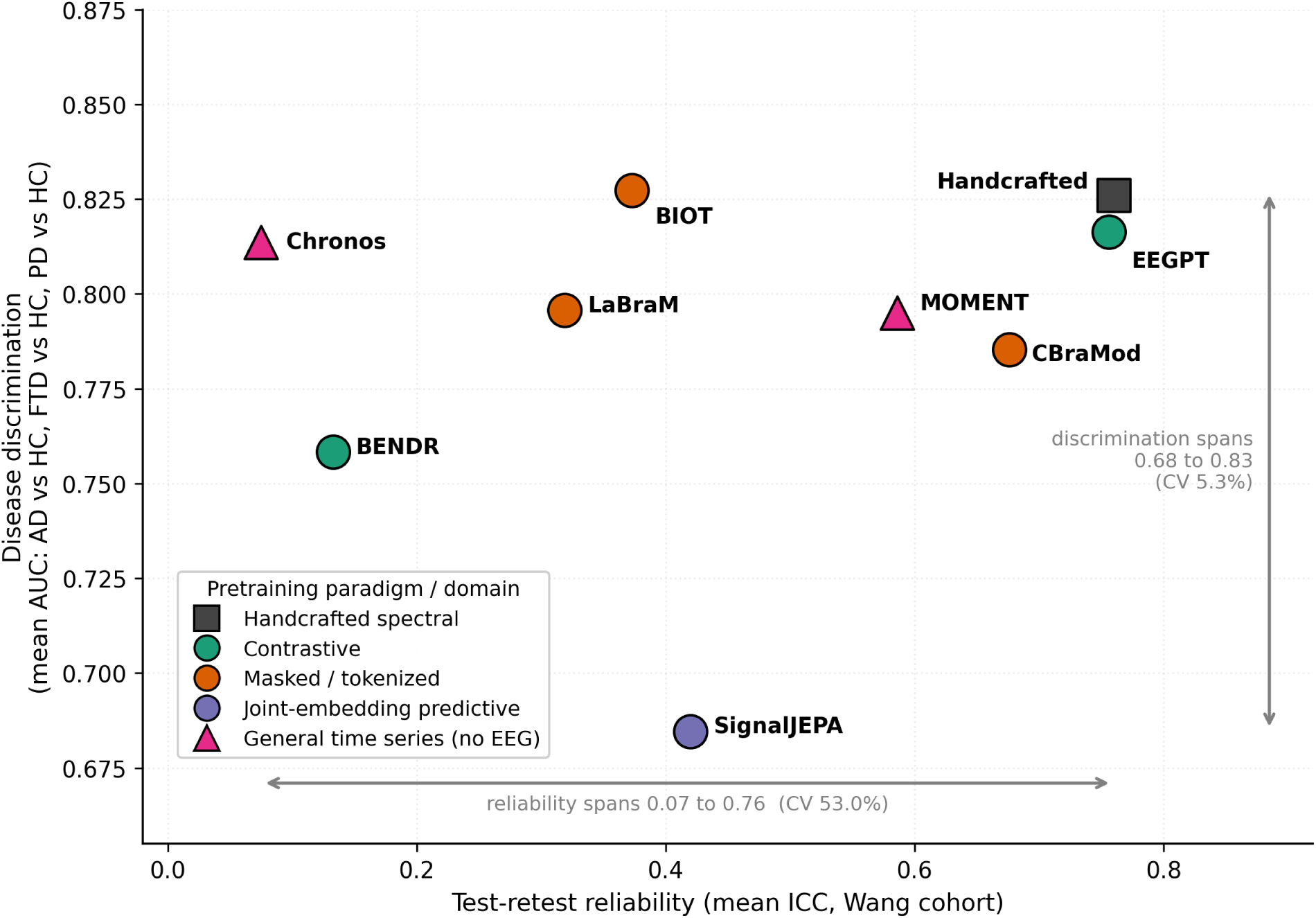
Reliability varies widely at comparable disease discrimination. Nine representations plotted by test-retest reliability (mean ICC, Wang cohort) against disease discrimination (mean AUC across the AD-versus-HC, FTD-versus-HC, and Cavanagh-PD contrasts), coloured and labelled by pretraining paradigm and domain. Points cluster vertically (discrimination AUC 0.69 to 0.83, coeffiicient of variation 5.4%) but spread across nearly the full horizontal range (reliability 0.08 to 0.76, coeffiicient of variation 53.0%). At comparable discrimination, reliability varies roughly tenfold more; the exact arithmetic is given in Methods 2.4.

#### Robustness analysis

Three perturbations were applied to each clean Wang recording before embedding, after which the full reliability analysis was re-run: electrode failure, simulated by marking 25% or 50% of channels bad and spline-interpolating them; duration truncation, keeping 50% or 25% of the original recording length; and additive Gaussian noise, scaled to 0.5 or 1.0 times the per-channel standard deviation. For each condition we computed mean per-dimension ICC and the Spearman rank correlation of the resulting ordering against baseline, and decomposed within- and between-subject variance to distinguish genuine reliability change from variance compression.

**Geometric mechanistic analyses**, run on three representative models spanning the reliability range (LaBraM, BENDR, and EEGPT), addressed two questions. The disease-discriminative subspace is the between-class subspace spanned by the globally centred class means of the disease cohort. With three classes that subspace has rank two, and it is constructed at that rank: an earlier version padded the two class-mean differences with principal directions of the same class-mean matrix to reach five dimensions, but those additional directions were numerically degenerate rather than disease-related, and comparing a rank-deficient subspace against a full-rank random null is not meaningful. The null is drawn at the dimension of the subject-stable subspace and compared with the same rank-two disease subspace, so observed and null overlaps average over the same number of principal angles. For recoverability, a linear adapter with output dimension 32 was fit on Wang embeddings using a supervised-contrastive objective on subject identity, trained on one split of subjects and evaluated by held-out test-retest ICC on a disjoint split, using five-fold cross-validation with a fixed seed. The adapter operated on the same mean-pooled recording-level embeddings used everywhere else in this paper, not on token-level or pre-pooling features. It therefore tests whether a subject-stable direction is recoverable by a linear map of the pooled representation, and cannot separate information lost in the encoder from information lost in pooling, a distinction Širca et al. (2026) show matters for these models. For subspace independence, we computed the principal-angle overlap, the mean cosine of principal angles, between the top between-subject directions in Wang (the subject-stable subspace), the top within-subject directions (the noise subspace), and the disease-discriminative directions in Miltiadous, which was defined on a disjoint disease cohort with no subjects shared with Wang. This was compared against a random-subspace null of matched dimension.

**Perturbational mechanistic analyses** used six representative models spanning the reliability range: EEGPT, CBraMod, LaBraM, BENDR, BIOT, and Chronos as a non-EEG-specific comparison. Frequency-band ablation applied zero-phase Butterworth band-stop filtering, defining delta as 1 to 4 Hz, theta as 4 to 8 Hz, alpha as 8 to 13 Hz, beta as 13 to 30 Hz, and gamma as 30 to 45 Hz, on the clean exported signal, then re-embedded and re-computed ICC. The outcome measure was the ICC drop, baseline minus ablated, with paired bootstrap confidence intervals. Layer-wise reliability used forward hooks on each model’s intermediate blocks, auto-discovered by module-name pattern and validated with a mandatory diagnostic pass that compared discovered layer count and shapes against each model’s known depth before any full extraction was run. Layer-wise pooling, taking the mean over sequence or time, is a generic heuristic that has not been independently verified against each model’s offiicial readout, so within-model depth trends should be trusted more than exact numerical agreement with a model’s published final-layer ICC. For AUC under ablation, band-ablated and layer-wise embeddings were extracted on the Miltiadous disease cohort, and AD-versus-HC and FTD-versus-HC discrimination AUC were computed with the identical methodology described above.

## 3. Results

The results are organised as an argument in five steps. We first establish that reliability varies widely and that this variation is neither predictable from conventional model descriptors nor attributable to dimensionality or acquisition artifacts (Section 3.1). We then ask what physiological information reliable representations encode (Section 3.2), and whether that information is the same information that supports disease discrimination (Section 3.3). Having found the two to be partially distinct, we ask where in a network this distinction is preserved or lost (Section 3.4), and finally which of these effects replicate at a longer retest interval (Section 3.5).

### 3.1 Reliability is not explained by conventional model descriptors

Using each subject’s longest available retest interval, the nine representations’ reliability spanned near-zero to high in both healthy cohorts (**Table 1**, **Figure 1**). Full values with bootstrap confidence intervals for both cohorts are in **Supplementary Table S6**. In Wang, at roughly one month and n = 58, handcrafted features (0.760), EEGPT (0.756), and CBraMod (0.676) were highly reliable. MOMENT (0.586, despite no EEG exposure), SignalJEPA (0.420), and BIOT (0.373) were intermediate. LaBraM (0.319) was poor, and BENDR (0.133) and Chronos (0.075, with a confidence interval crossing zero) were the lowest of all nine. In Henao, at roughly two years and n = 34 to 39, handcrafted (0.653), CBraMod (0.663), and EEGPT (0.609) remained the reliable cluster. LaBraM (0.533), BIOT (0.555), and Chronos (0.571) rose substantially from their Wang values, while BENDR (0.152) remained the lowest.

**Table 1.** Reliability, distinctiveness, and disease discrimination for nine representations. Reliability is the mean per-dimension ICC(2,1), computed on each subject’s longest-interval pair and averaged across embedding dimensions. The agreement ratio is the multivariate within/between distance ratio under the shift-sensitive distance, the counterpart of the absolute-agreement ICC (Methods 2.4); lower values indicate greater reliability. The consistency-distance variant and the Henao Isaza values are in **Supplementary Table S7**. Identifiability chance level is 0.011. AD and FTD AUC are from permutation-validated repeated stratified 5-fold cross-validation, significant at p < 0.05 for all nine representations except SignalJEPA on the PD contrasts and for the Rockhill-PD cohort generally (**Supplementary Table S5**).

| Representation<br>(paradigm) | ICC<br>Wang | ICC<br>Henao | Agreement<br>ratio<br>(Wang) | Identifiability | AUC<br>AD vs<br>HC | AUC<br>FTD<br>vs HC |
| --- | --- | --- | --- | --- | --- | --- |
| Handcrafted | 0.760 | 0.653 | 0.450 | 0.229 | 0.849 | 0.834 |
| EEGPT (contrastive) | 0.756 | 0.609 | 0.460 | 0.711 | 0.876 | 0.770 |
| CBraMod (masked) | 0.676 | 0.663 | 0.544 | 0.602 | 0.854 | 0.778 |
| MOMENT (general<br>time series) | 0.586 | 0.696 | 0.525 | 0.500 | 0.801 | 0.763 |
| SignalJEPA<br>(predictive) | 0.420 | 0.334 | 0.765 | 0.120 | 0.884 | 0.790 |
| BIOT<br>(masked/tokenized) | 0.373 | 0.555 | 0.603 | 0.572 | 0.858 | 0.798 |
| LaBraM (masked) | 0.319 | 0.533 | 0.639 | 0.633 | 0.899 | 0.792 |
| BENDR (contrastive) | 0.133 | 0.152 | 0.946 | 0.018 | 0.790 | 0.694 |
| Chronos (general time<br>series) | 0.075 | 0.571 | 0.683 | 0.422 | 0.859 | 0.773 |

Quantified against discrimination, the contrast is stark. Across these same nine representations, reliability has a coeffiicient of variation of 53.0% (mean ICC 0.455 in Wang), whereas mean diseasediscrimination AUC has a coeffiicient of variation of 5.4% (mean 0.789 across the three adequately powered contrasts). Reliability therefore varies roughly tenfold more than discrimination does across the representations tested, a ratio of 9.9 (**Figure 1**; the arithmetic and a sensitivity check are given in Methods 2.4). We did not test formal AUC equivalence, so we describe discrimination as varying substantially less than reliability rather than as equivalent across representations.

This variation tracked neither pretraining paradigm nor pretraining domain. The two masked-reconstruction models fell at opposite ends, with CBraMod reliable and LaBraM not. The two contrastive models did too, with EEGPT reliable and BENDR the least reliable of all nine in Wang. Within-paradigm variation was as large as between-paradigm variation. Pretraining domain fared no better as a predictor. MOMENT, with zero EEG exposure, was among the more reliable representations in both cohorts, and the single highest of all nine in Henao. Chronos, which shares MOMENT’s non-EEG pretraining domain but uses a different architecture, was the least reliable representation of all nine in Wang, the largest cross-cohort reversal of any representation in this study and a pattern we return to in Section 3.5.

This ordering is real, not a dimensionality or acquisition artifact. Mean ICC averages over dimensionalities ranging from 13 to 2048, so we also computed dimensionality-agnostic ratios of within-subject to between-subject representational distance. The agreement ratio, the variant that penalises session shifts as ICC(2,1) does, reproduced the ICC ordering in both cohorts (Spearman rho = -0.85, p = 0.004 in Wang; rho = -0.75, p = 0.020 in Henao Isaza; **Table 1**, **Supplementary Table S7**): handcrafted and EEGPT were lowest, meaning most reliable, and BENDR was highest, meaning least reliable, approaching a value of 1.0 where an individual’s repeated recordings are nearly as far apart as different individuals’ recordings. The consistency ratio, which uses correlation distance and is blind to such shifts, agrees with the ICC ordering in Wang (rho = -0.90, p = 0.001) but not in Henao Isaza (rho = -0.45, p = 0.224), where CBraMod is second most reliable by ICC and eighth of nine by that ratio. We read this as a property of the metric rather than of the models: correlation distance discards exactly the session-to-session offsets that an absolute-agreement ICC counts, and the discrepancy disappears when the distance is matched to the ICC definition. It is also a caution that a consistency-type multivariate measure should not be used to corroborate an absolute-agreement one. A matched-dimensionality PCA control confirmed that the reliable models remained above LaBraM, and BENDR remained lowest, across informative variance ranges (**Supplementary Tables S7 and S8**, **Supplementary Figure S1**). Subject identifiability, at a chance level of 0.011, dissociated informatively from stability. LaBraM is highly distinctive (0.633) yet unstable (ICC 0.319): recognisable, but drifting. BENDR is neither distinctive nor stable, while EEGPT and MOMENT are both.

The ordering also survived realistic acquisition degradation (**Figure 2b**). This analysis was run on the 54 Wang subjects for whom all seven perturbation conditions could be regenerated, so its baseline values differ slightly from the n = 58 values in **Table 1**; comparisons below are made within the stress-test set. Across electrode failure, with Spearman rho of 1.00 at 25% of channels lost and 0.96 at 50%, and shortened recordings, with rho of 0.96 at 50% duration and 0.89 at 25%, the reliability rank order barely changed (**Figure 2c**; per-condition values in **Supplementary Table S9**). Handcrafted features and EEGPT remained the two most reliable representations in every condition, spanning 0.731 to 0.773 and 0.725 to 0.764 respectively, and BENDR remained the least reliable in every condition. Heavy additive noise partly broke the ordering, with rho of 0.86 at 0.5 times the per-channel signal standard deviation and 0.75 at 1.0 times, but variance decomposition showed this was compression rather than genuine improvement. LaBraM’s apparent ICC roughly doubled under noise, from 0.308 at baseline to 0.619 at 0.5 times and 0.632 at 1.0 times, while its within- and between-subject standard deviations both collapsed together, by 82% and 72% respectively, rather than reliability genuinely improving (**Supplementary Table S10**). Handcrafted features and EEGPT held both variance components stable under noise, with changes within plus or minus 10%, the diagnostic signature of a representation that is reliable for the right reason rather than as an artifact of variance compression. This robustness analysis was run for the seven EEG-oriented representations; MOMENT and Chronos were not yet included, a gap noted in Limitations.

**Figure 2.**
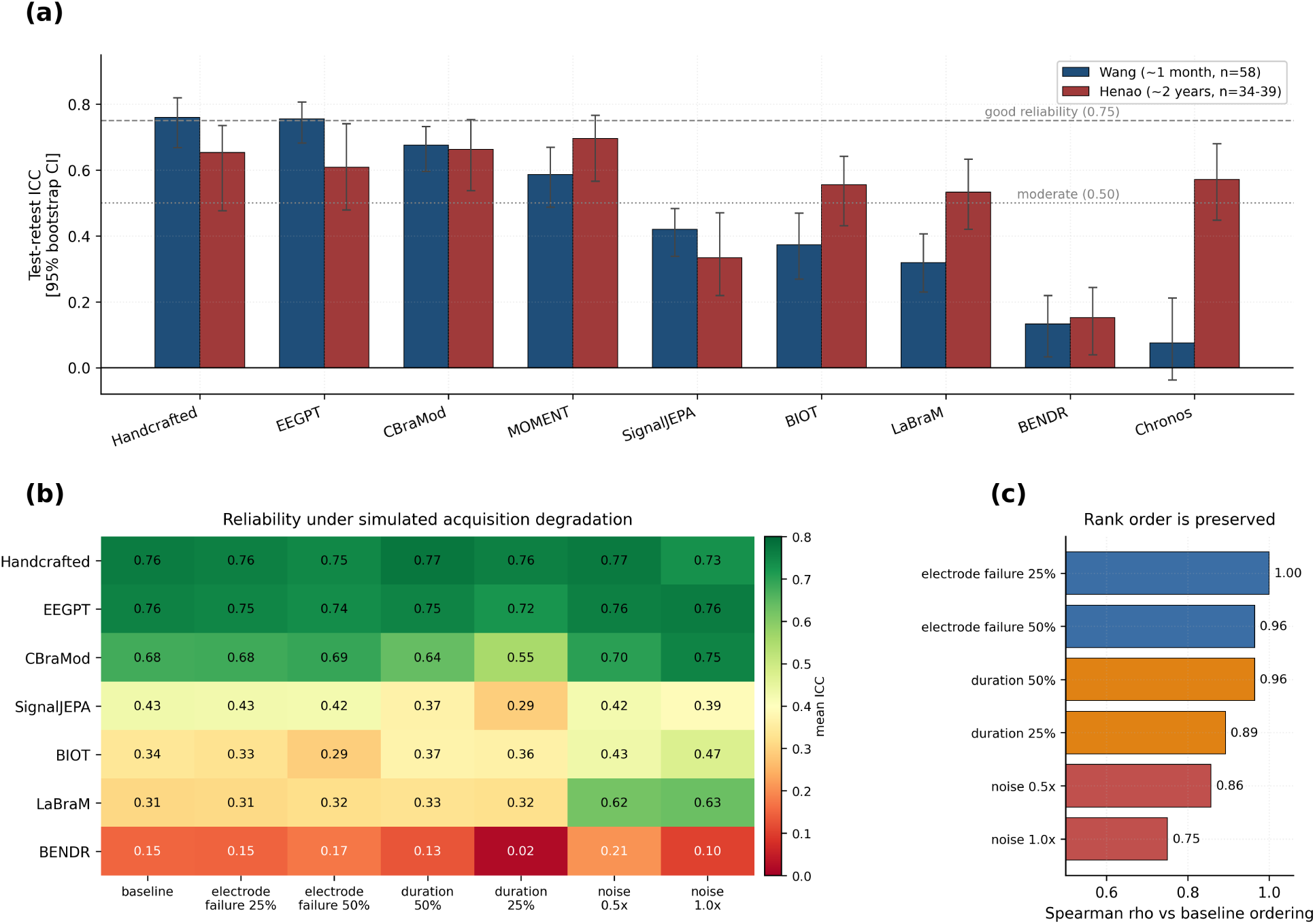
Reliability is a property of the trained representation, not an artifact of dimensionality or acquisition quality. (a) Mean ICC(2,1) per representation in the Wang (approximately one month) and Henao (approximately two years) retest cohorts, with 95% bootstrap confidence intervals, ordered by Wang reliability. Reference lines mark conventional thresholds for moderate (0.50) and good (0.75) reliability. The reliable cluster of handcrafted features, EEGPT, and CBraMod, and BENDR’s position at the bottom, are consistent across both intervals. (b) Acquisition-degradation stress test in Wang, on the 54 subjects for whom all seven conditions could be regenerated. Reliability was recomputed after simulated electrode failure, recording truncation, and additive Gaussian noise. (c) Spearman rank correlation between each degraded condition’s reliability ordering and the baseline ordering. Rank order is essentially preserved under electrode failure and truncation (rho 0.89 to 1.00) and partly breaks only under heavy noise, where variance decomposition shows the apparent gain in LaBraM to be variance compression rather than improved reliability.

Reliability differences are also not simply a byproduct of one fixed preprocessing choice. We compared handcrafted features and LaBraM under a minimal-preprocessing pipeline, using only band-pass filtering, average referencing, and unit normalisation, without the full pipeline’s bad-channel interpolation and epoch rejection. The unified pipeline substantially improved handcrafted-feature reliability, from 0.522 to 0.760, while if anything reducing LaBraM’s, from 0.449 to 0.319 (**Supplementary Table S11**). We do not isolate which preprocessing step drives either change, and we did not repeat this comparison for the other representations, so we do not claim the pattern generalises. What it establishes is narrower but still useful: a preprocessing effect on reliability can be representation-specific rather than uniform, while the ordering between the two representations tested here is preserved at both preprocessing levels.

### 3.2 Reliable representations encode specific physiological information

Variance decomposition shows that reliable representations concentrate embedding variance in the between-subject component rather than in session-to-session noise (**Supplementary Figure S2**), with correspondingly short session-to-session displacement in embedding space (**Supplementary Figure S3**). This is independently confirmed by linear CKA (Kornblith et al., 2019), a similarity measure constructed in an entirely different way from ICC (**Supplementary Figure S4**). Correlating each embedding dimension against the 13 handcrafted features shows that for CBraMod, EEGPT, and BIOT, the most reliable dimensions correlate most strongly with absolute alpha power (**Figure 3a**). For LaBraM the strongest correlate is alpha peak frequency instead, still an alpha-band feature, just not absolute power specifically. The test-retest reliability and between-subject variability of the 13 handcrafted features themselves, which the following two paragraphs draw on, are shown in **Figure 3b**.

**Figure 3.**
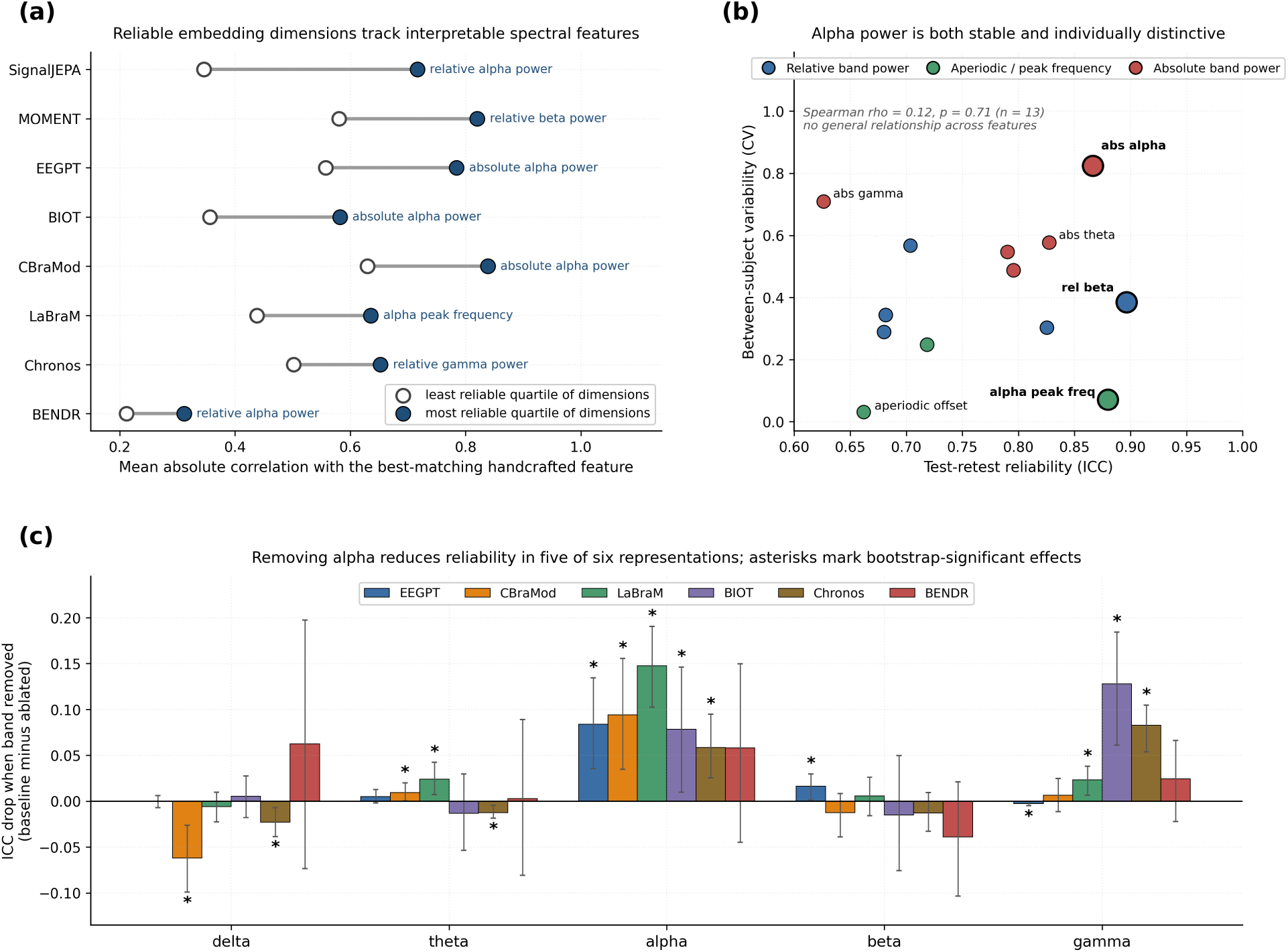
Alpha-band information is a candidate spectral contributor to reliability. (a) Dimension-physiology association in the Wang cohort. For each representation, embedding dimensions were ranked by their own test-retest ICC and split into quartiles. Open circles give the mean absolute correlation with the best-matching handcrafted feature for the least reliable quartile, filled circles the same quantity for the most reliable quartile, and the connecting bar the difference. In all eight representations the most reliable dimensions correlate more strongly with an interpretable spectral feature than the least reliable ones do. The annotation names the feature most often matched in the top quartile, which is an alpha-band feature in six of eight cases. (b) The stability-by-individuality pattern across all 13 handcrafted features. Absolute alpha power combines high test-retest reliability with the highest between-subject variability of the five power bands, whereas relative beta power is comparably reliable but among the least variable between subjects. Alpha peak frequency is the counterexample that prevents this being read as a general law: it is highly reliable yet almost invariant between subjects. Across all 13 features the relationship is not significant (Spearman rho = 0.12, p = 0.71), so this is an alpha-specific observation rather than a general rule. (c) Frequency-band ablation in Wang. Bars show the ICC drop, baseline minus ablated, when each band is removed by zero-phase band-stop filtering before re-embedding, with 95% bootstrap confidence intervals from 1000 draws. Positive values indicate that removing the band reduced reliability. Asterisks mark effects whose confidence interval excludes zero. Alpha is significant in five of the six representations tested, and BENDR is the exception, showing no significant effect for any band.

Frequency-band ablation supports the same conclusion through a different route. Removing alpha produced a reproducible, bootstrap-confirmed reduction in reliability in five of six representations tested on Wang (**Figure 3c**): EEGPT (+0.084, 95% CI +0.035 to +0.134), CBraMod (+0.094, +0.035 to +0.156), LaBraM (+0.148, +0.103 to +0.191), BIOT (+0.078, +0.010 to +0.146), and Chronos (+0.058, +0.025 to +0.095). Alpha was the largest single-band effect in three of these five representations, EEGPT, CBraMod, and LaBraM. In the remaining two, BIOT and Chronos, gamma was marginally larger, with alpha a close second in both cases (the full band-by-band break-down is in **Supplementary Table S12**). BENDR showed no significant band-ablation effect at all, consistent with its low reliability being concentrated in a specific architectural failure discussed in Section 3.4, rather than distributed dependence on any one band. Taken together, this correlational and perturbational evidence identifies alpha-band information as a candidate spectral contributor to reliability rather than establishing where reliability originates in a comprehensive or exclusive sense.

This is not a trivial consequence of alpha simply being the most reliable raw-signal band. In the handcrafted-feature ICCs, relative beta power (0.897) is in fact the single most reliable of all 13 features, exceeding absolute alpha power (0.867), and average raw-signal reliability ranks alpha and beta identically at 0.846 each. Yet beta’s band-ablation importance is negligible in every model tested, with the largest ICC drop only +0.016, while alpha’s is the largest or second-largest effect throughout. A better-supported explanation combines two properties: alpha has high raw-signal reliability and high between-subject variability, with a coeffiicient of variation of 0.82, the highest of the five power bands, while beta has comparable reliability but the lowest between-subject variability, at 0.49. Alpha is stable within a subject and distinctive between subjects. Beta is stable but comparatively generic (**Figure 3b**).

This “stability times individuality” pattern does not generalise. Tested across all 13 handcrafted features, ICC and between-subject CV show no significant relationship (Spearman rho = 0.12, p = 0.71, n = 13). Alpha peak frequency is the second most reliable feature of all 13, with ICC = 0.880, consistent with its established status as a trait-like marker that resists even sustained cognitive intervention (Grandy et al., 2013), yet it has the second-lowest between-subject variability of any feature, with CV = 0.07, behind only the aperiodic offset’s CV of 0.03. That is the opposite combination the hypothesis predicts, and the feature is still highly reliable. The calibrated claim is this: alpha power specifically combines high reliability with high between-subject variability, and this correlates with its outsized influence on model reliability, but “stable and individually distinctive” is not a general property that separates reliable from unreliable EEG features overall. Alpha should be understood as one identified contributor to reliability, not a complete account of it.

Gamma adds a further complication. It is the second-largest band-ablation driver of reliability, with a mean drop of 0.044 across all six perturbation-tested representations, despite having the lowest raw-signal reliability of any band, averaging 0.665 across its absolute and relative power features (**Supplementary Figure S5**). Unlike alpha, it is nearly irrelevant to disease discrimination, with a mean AUC drop of just 0.007 (Section 3.3). One candidate explanation is EMG or muscle-artifact contamination. Two crude but standard proxies, the frontotemporal-to-central gamma-power ratio and gamma-band kurtosis, show moderate test-retest reliability of 0.500 (95% CI 0.269 to 0.747) and 0.404 (0.161 to 0.627) respectively in Wang, n = 58, comparable to and statistically indistinguishable from the ICC of absolute gamma power itself (0.626) through overlapping confidence intervals. This is consistent with, though not proof of, the artifact hypothesis; an ICA-based ground-truth decomposition would be needed to make the claim stronger.

### 3.3 Reliability and disease discrimination are partially dissociable

This is the paper’s central finding, and we report it through two conceptually and methodologically distinct analyses. Both operate on representations derived from the same EEG signals, but they were developed independently, probe different aspects of the embedding space, and converge on the same conclusion.

#### Geometric evidence, from subspace geometry

Two analyses of the embedding space, both independent of the frequency-domain evidence below, bear on whether the subject-stable and disease-discriminative structure is the same structure. First, a supervised-contrastive linear adapter fit on Wang embeddings substantially recovered LaBraM’s reliability, from a held-out ICC of 0.29 to 0.49, indicating that a subject-stable subspace exists within LaBraM’s frozen features but is not exposed by the default representation (**Figure 4a**; per-fold values in **Supplementary Table S14**). The identical procedure did not recover BENDR, which moved from 0.12 to 0.08; we do not read this as proof that no stable subspace exists anywhere in BENDR’s features, only that none was detected by this particular approach. Second, using principal angles, we measured the overlap between the top between-subject directions in Wang healthy embeddings, the subject-stable subspace, and the disease-discriminative directions in Miltiadous, defined independently with no shared subjects, for LaBraM, BENDR, and EEGPT. The disease subspace is the between-class subspace of the three class means, which has rank two, and the random null is drawn to compare across the same number of principal angles (Methods 2.4). Across all three representations the overlap fell at or below that null; for LaBraM the observed overlap was 0.187 against a null 95th percentile of 0.191, and the stable subspace carried no more disease information than the within-subject noise subspace did, which for BENDR and LaBraM carried slightly more (**Figure 4b**; **Supplementary Table S13**). No above-null overlap was detected in any model tested. This is an absence of detected overlap rather than a demonstration of independence, and LaBraM sits close to its threshold, at 0.187 against 0.191; a formal independence claim would need an equivalence margin, which we do not attempt.

**Figure 4.**
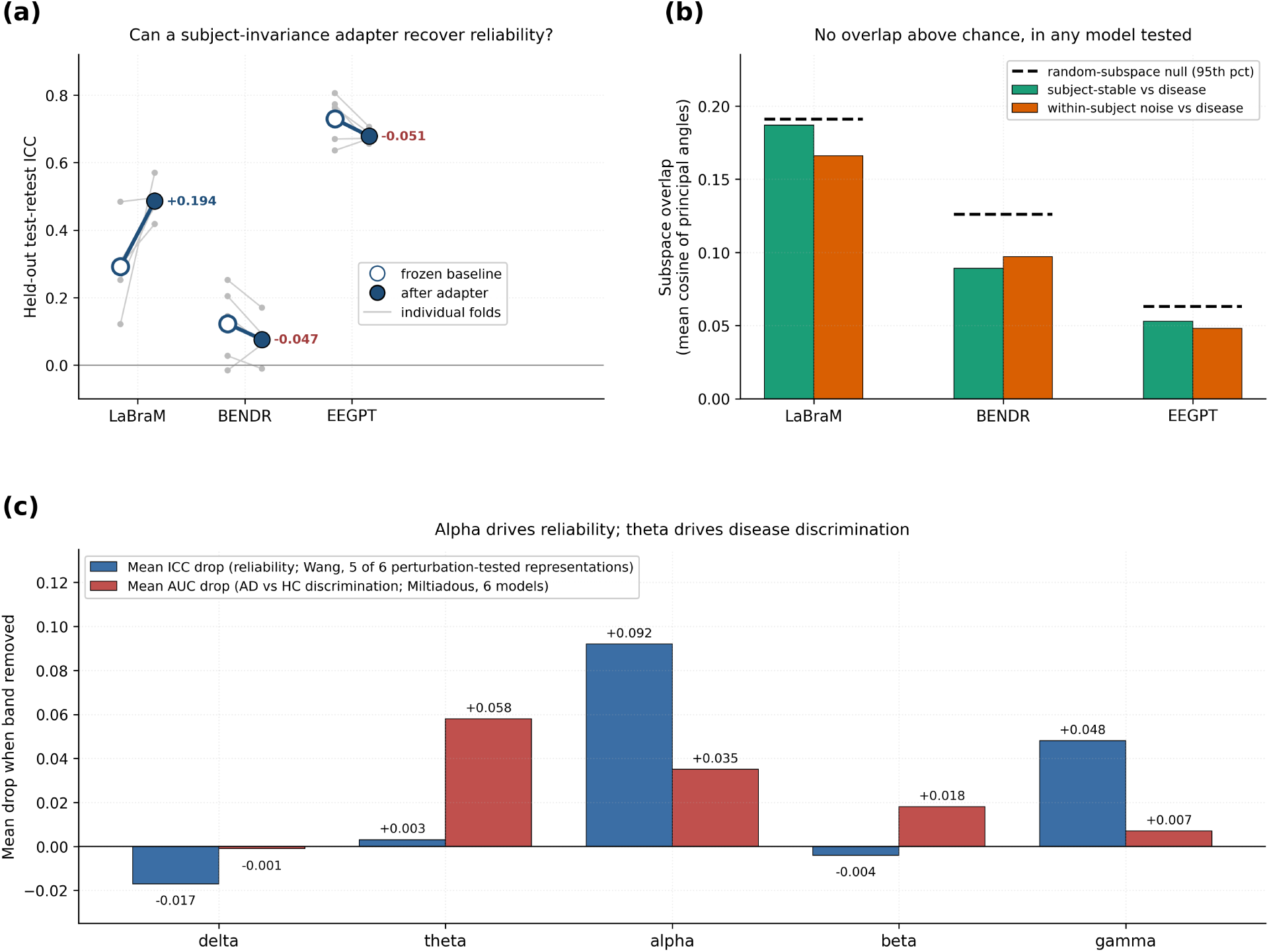
Reliability and disease discrimination draw on partially distinct information, by two independent lines of evidence. (a) Geometric evidence, recoverability. A supervised-contrastive linear adapter was fit on Wang embeddings and evaluated on a disjoint split of subjects. Open circles give the frozen baseline, filled circles the value after adaptation, faint lines the five individual cross-validation folds, and the annotation the mean change. LaBraM’s held-out reliability rises from 0.29 to 0.49, BENDR shows no gain, and the already-reliable EEGPT is mildly compressed. (b) Geometric evidence, subspace independence. Principal-angle overlap between the subject-stable subspace defined in Wang and the disease-discriminative subspace defined independently in Miltiadous, with the within-subject noise subspace shown for comparison. Dashed lines mark the 95th percentile of a random-subspace null of matched dimension. In all three models the observed overlap falls at or below that null and does not exceed the noise-subspace overlap. (c) Spectral evidence. Mean drop in reliability (Wang) and in AD-versus-HC discrimination (Miltiadous) when each band is removed. Alpha shows the largest reliability drop and theta the largest discrimination drop, so the band that matters most for one property is not the band that matters most for the other. Values correspond to **Table 2**.

**Table 2.**
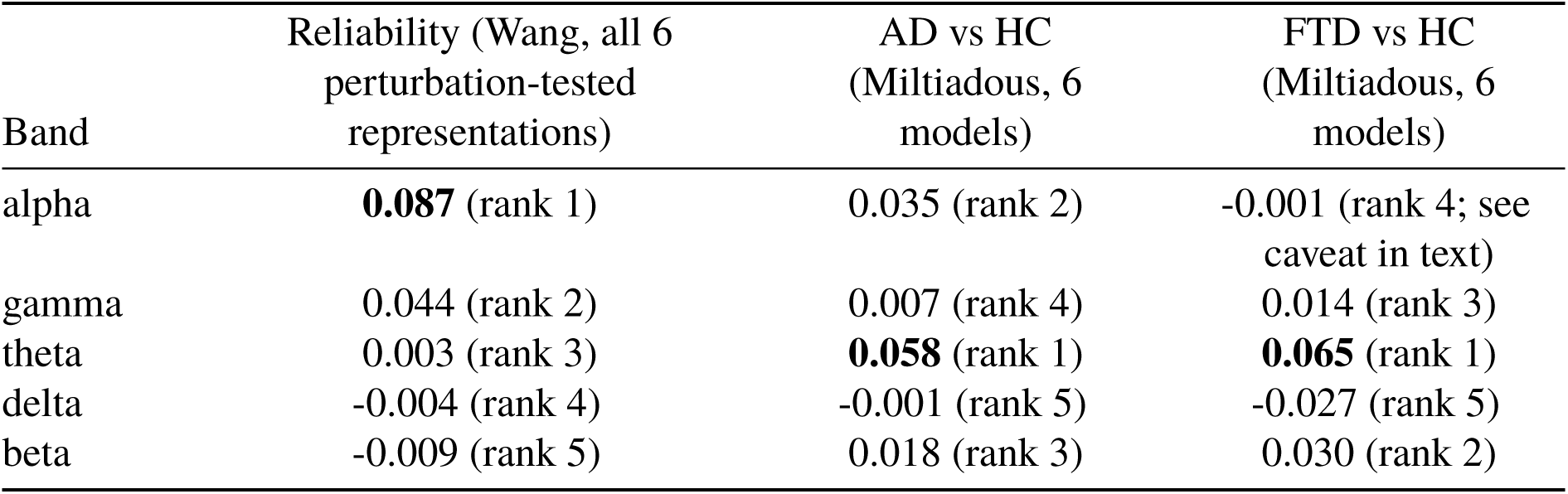
Mean AUC/ICC drop when each band is removed, by outcome. The reliability column is the mean over all six perturbation-tested representations, namely EEGPT, CBraMod, LaBraM, BIOT, BENDR and Chronos. All six are pooled irrespective of whether their individual band effects reached significance, so that no representation is dropped on the basis of its own result. As a sensitivity check, omitting BENDR, whose effects are uniformly non-significant, gives 0.092 for alpha and 0.048 for gamma, which remain ranks one and two. Note that Chronos is included here and is not EEG-pretrained. AD-versus-HC uses permutation testing only for the baseline, no-alpha, and no-theta conditions; FTD-versus-HC has full five-band permutation testing for all six models. Those permutation tests shuffle diagnostic labels and therefore establish that each ablated representation still discriminates above chance, surviving within-family BH-FDR correction with the exceptions detailed in Methods 2.4; they do not test whether removing a band changed the AUC. That change is tested separately by paired bootstrap in **Supplementary Table S17**, where the ranking reproduces but no individual comparison survives correction. Ranks are within each column. These values are plotted in **Figure 4c**.

| Band | Reliability (Wang, all 6<br>perturbation-tested<br>representations) | AD vs HC<br>(Miltiadous, 6<br>models) | FTD vs HC<br>(Miltiadous, 6<br>models) |
| --- | --- | --- | --- |
| alpha | **0.087** (rank 1) | 0.035 (rank 2) | -0.001 (rank 4; see<br>caveat in text) |
| gamma | 0.044 (rank 2) | 0.007 (rank 4) | 0.014 (rank 3) |
| theta | 0.003 (rank 3) | **0.058** (rank 1) | **0.065** (rank 1) |
| delta | -0.004 (rank 4) | -0.001 (rank 5) | -0.027 (rank 5) |
| beta | -0.009 (rank 5) | 0.018 (rank 3) | 0.030 (rank 2) |

#### Spectral evidence, from band ablation

Using the Miltiadous AD, FTD, and HC cohort with the same band-ablation manipulation, reliability and AD-versus-HC discrimination are driven by substantially different bands (**Table 2**). Theta is the largest driver of AD-versus-HC discrimination, with a mean AUC drop of 0.058, despite being nearly irrelevant to reliability, with a mean ICC drop of only 0.003. The two sides of that comparison are not equally well supported, and we tested them to the same standard rather than leaving the asymmetry implicit. A paired subject-level bootstrap on the AUC drop itself, the direct analogue of the interval already reported for the ICC drop, confirms the ranking in both disease contrasts, with theta first and a mean drop of 0.061, but no individual model-by-band comparison survives Benjamini-Hochberg correction within either contrast, and only eight of sixty have confidence intervals excluding zero uncorrected, five of them theta (**Supplementary Table S17**). By the same uncorrected standard the reliability side is markedly stronger: alpha’s ICC drop excludes zero in five of six representations. The band ranking is therefore a reproducible point-estimate pattern on both axes, with direct statistical support for the reliability half and, at this sample size, only suggestive support for the discrimination half. This is directionally consistent with established EEG evidence: theta-band slowing is a recognised correlate of Alzheimer’s disease and cognitive decline (Musaeus et al., 2018), and theta-based spectral power ratios discriminate frontotemporal dementia in the same public cohort we use here (Chang and Chang, 2023). This is a link we did not set out to confirm, but it lends construct validity. Alpha contributes to both axes, ranking first for reliability and second for AD-versus-HC discrimination, so this is not a fully clean, disjoint decoupling. The precise claim is narrower: the band most important for reliability, alpha, is not the band most important for discrimination, theta.

A second disease contrast generalises the pattern, and sharpens it. Extending full permutation testing to FTD-versus-HC (n = 22 FTD, 29 HC), theta remains the largest driver of discrimination, at 0.065, rank one (**Table 2**, FTD-versus-HC column), and five of six representations show small negative alpha drops ranging from -0.005 to -0.084, consistent with alpha being unimportant to FTD discrimination just as it was for AD. All six models’ FTD-versus-HC baselines are individually significant, at p ≤ 0.04 and five of six at p ≤ 0.025. One caveat applies to the reported mean: BENDR is an outlier with a large positive alpha-removal drop of +0.230, and its no-alpha AUC is not even above chance (p = 0.68). The table’s near-zero mean of -0.001 therefore partly reflects this single outlier cancelling the other five models’ small negative values, rather than six models independently showing no alpha effect. The five-of-six pattern, not the pooled mean, is the more informative summary.

#### A third, architecture-internal demonstration

Layer-wise AUC probing on Miltiadous shows that LaBraM’s AD-versus-HC discrimination peaks at block 6, with AUC = 0.934, and drops notably by block 8, to AUC = 0.898, the third-lowest of 12 layers, the same block that is LaBraM’s reliability peak (ICC = 0.484, Section 3.4). The two profiles are plotted on a shared layer axis in **Figure 5c** and **Figure 5d**. We tested this pattern’s statistical robustness directly with a paired bootstrap of 500 draws on the AUC difference between layers. Block 6 versus block 8 gave +0.013 (95% CI -0.012 to +0.040); block 6 versus the published final layer, block 11, gave +0.006 (-0.032 to +0.043); and block 8 versus final gave -0.006 (-0.041 to +0.028). All three confidence intervals include zero, so none is significant at this sample size of n = 63, and none is corrected for multiple comparisons (Methods 2.4). The reliability side of this crossover, block 8 versus the final layer, is solidly supported by non-overlapping confidence intervals. The discrimination side is a real, visually clear point-estimate pattern that is not yet confirmed as statistically significant.

**Figure 5.**
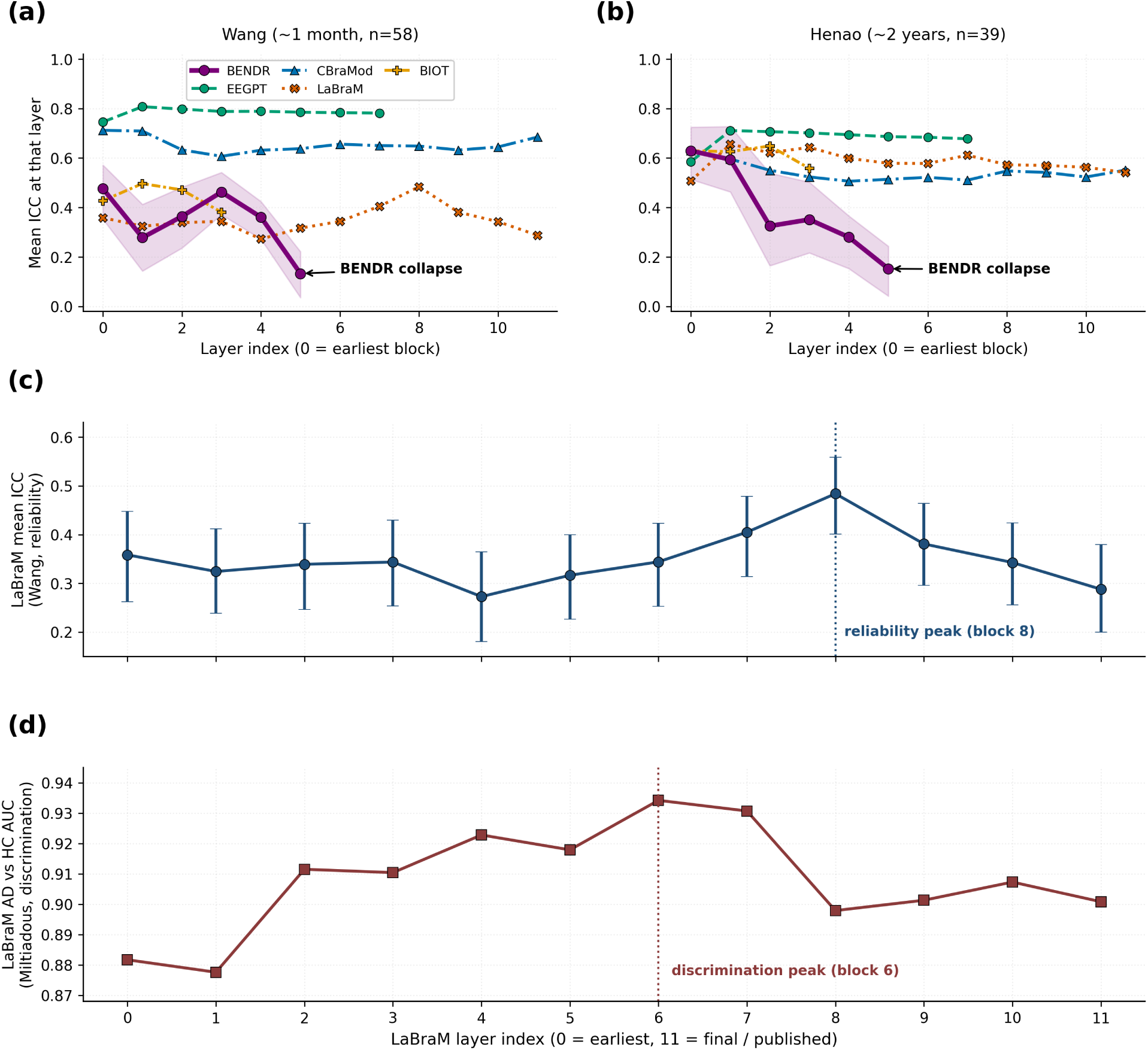
Network depth shapes whether reliability and discrimination are preserved, traded off, or jointly degraded. (a) Layer-wise mean ICC in the Wang cohort. BENDR is drawn in black with its 95% bootstrap confidence band shaded; it falls from 0.476 at layer 0 to 0.133 at its final encoder stage, a non-overlapping drop. EEGPT and CBraMod hold reliability approximately constant across their full depth. (b) The same analysis in Henao, where BENDR’s collapse replicates at an independent retest interval, from 0.629 to 0.152. Series in A and B are distinguished by marker shape and line style as well as tone, so they remain separable in greyscale. (c) LaBraM’s layer-wise reliability in Wang, with 95% bootstrap confidence intervals from 1000 draws. (d) LaBraM’s AD-versus-HC discrimination AUC in Miltiadous, on the same layer axis as C. Plotting the two measures as separate panels on a shared axis, rather than on two vertical axes of one panel, avoids implying a direct numerical comparison between an ICC and an AUC. Reliability peaks at block 8 and discrimination at block 6, so no single layer is simultaneously optimal for both. The reliability side of this crossover is supported by non-overlapping confidence intervals; the discrimination side is a point-estimate pattern that the paired bootstrap does not support at this sample size (Section 3.3).

Combined, three analyses provide convergent, though uneven, evidence for the same conclusion using three distinct methodologies: subspace geometry, derived from embedding-space directions; spectral band ablation, derived from perturbing the input signal; and architecture-internal layer crossover, derived from network depth. Reliability and disease discrimination draw on partially, not fully, independent information.

This dissociation also holds at the level of whole representations, not just bands or layers. All nine representations achieved broadly similar disease discrimination, with AD AUC from 0.79 to 0.90 and FTD from 0.69 to 0.83, all significant except where noted (**Table 1**; full results in **Supplementary Table S5**), and with no consistent leader. Parkinson’s discrimination was markedly more cohort-dependent: eight of nine representations reached significance in Cavanagh, at a mean AUC of 0.74, whereas only one did so in the smaller Rockhill cohort, where the mean AUC was 0.47. AD and FTD discrimination is therefore broadly preserved across representations, while PD discrimination is not, and we treat only the former as the well-powered comparison. Reliable and unreliable models achieved overlapping AUCs, most starkly for BENDR, which discriminated AD and PD as well as other models, at AUC 0.79, while being the least reliable representation by every metric. The exception was SignalJEPA, which matched the others on AD and FTD but failed on both PD cohorts. MOMENT and Chronos, despite no EEG exposure, discriminated disease comparably to the EEG-oriented representations, with MOMENT second of nine on PD-Cavanagh at 0.821, behind BIOT at 0.826. Formally, the representation-level rank correlation between reliability and mean discrimination AUC across the three adequately powered contrasts was weak and not distinguishable from zero (Spearman rho = 0.233, p = 0.546, n = 9), corroborating the band- and layer-level dissociation above at the level of representations as a whole.

Taken together, the evidence in this section converges, and it converges on a partial separation rather than a clean one. Four analyses bear on the same question by different routes: a recoverability probe and a subspace-overlap test that examine the geometry of the embedding space, a band-ablation experiment that manipulates the input signal instead, and a representation-level rank correlation that uses neither. All four indicate that the information supporting subject stability is not the information supporting disease discrimination. The convergence carries weight because the approaches fail in different ways, so a shared methodological artifact is an unlikely explanation for all of them at once. The separation is nonetheless incomplete: alpha remains the second-ranked band for AD-versus-HC discrimination, and the subspace overlaps sit at rather than below the random-subspace null, so the two properties draw on overlapping but non-identical information. The section that follows asks whether this partial separation survives passage through a network’s depth.

### 3.4 Network depth shapes how these properties are preserved

Layer-wise reliability probing in the Wang cohort shows that models differ qualitatively, not just quantitatively, in how reliability is distributed across depth, and this pattern is itself architecture-specific (**Figure 5**). The clearest single result is architectural: network depth can destroy measurement reliability outright, and the effect is reproducible.

BENDR is the clearest case. Its reliability collapses at the final encoder stage, and AUC collapses in the same direction, making this a pattern of shared destruction rather than a tradeoff. At layer 0, ICC = 0.476 (95% CI 0.353 to 0.572); at the final layer, layer 5, ICC = 0.133 (0.035 to 0.221) (**Figure 5a**). A paired subject-level bootstrap over the same 58 subjects puts the drop at 0.339 (95% CI 0.205 to 0.479), p ≤ 0.002 at the resolution of 1000 draws, and it survives Benjamini-Hochberg correction within the layer-wise reliability family. This replicates cleanly at Henao, where layer 0 is 0.629 (0.514 to 0.725) and the final layer 0.152 (0.042 to 0.243), a paired drop of 0.478 (0.330 to 0.621) over 39 subjects, also surviving correction (**Figure 5b**). Confirmed at two independent retest intervals, it is the single most robust layer-wise finding in this study. The recoverability probe reported in Section 3.3 bears on this directly: a linear adapter recovered LaBraM’s reliability but not BENDR’s, which indicates that BENDR’s final-stage representation does not retain subject-stable information in a form recoverable by the evaluated linear adapter. This does not distinguish information that has been destroyed from information that is present but inaccessible to a linear readout (**Figure 4a**).

EEGPT and CBraMod behave in the opposite way. Both establish reliability early and hold it stable through depth, with EEGPT ranging from 0.75 to 0.81 across all eight blocks and CBraMod from 0.61 to 0.71 across all 12 layers. Layer-wise AUC shows no strong tradeoff in either model, so both properties appear jointly achievable in these two architectures. This matters for interpreting BENDR: its collapse is a property of that architecture, not an inevitable consequence of depth.

LaBraM occupies an intermediate position, and the evidence for it is weaker and more exploratory than for BENDR. Its reliability is non-monotonic with depth and does not peak at the default final layer. At block 8 of 12, ICC = 0.484 (0.401 to 0.558), against the published final block’s 0.288 (0.200 to 0.379); the paired difference is 0.192 (0.147 to 0.234), p ≤ 0.002, and survives correction within the same family (**Figure 5c**). Its discrimination peaks at a different depth still, at block 6 (Section 3.3, **Figure 5d**), so no single LaBraM layer is simultaneously optimal for both properties. Unlike the reliability side, the AUC side of this pattern is not statistically supported: the paired bootstrap difference between blocks 6 and 8 is 0.013 (95% CI -0.012 to 0.040), p = 0.40, and none of the three pairwise layer contrasts survives correction, the smallest corrected value being 0.84. BIOT, with only four layers available, shows a weak echo of the same pattern, though it was not formally significance-tested layer to layer given how few layers exist. Chronos shows a depth profile that inverts between cohorts, which we discuss in Section 3.5 alongside the cross-cohort reliability anomaly it forms part of.

Taken together, whether a model’s depth trades reliability against discrimination, preserves both, or degrades both jointly is an architecture-specific property rather than a general rule, a typology this study did not set out to find and that may be of independent interest for EEG-FM architecture design.

### 3.5 Longitudinal replication reveals conserved and architecture-specific effects

At the longer Henao interval the alpha-ablation reliability effect replicates in four of the six perturbation-tested representations, CBraMod, EEGPT, LaBraM and Chronos, whose bootstrap confidence intervals exclude zero (**Figure 6a**). Restricting attention to the five EEG-pretrained models, three replicate. Chronos, which has no EEG exposure, is among those that do, which is consistent with the wider pattern that EEG-specific pretraining is not what distinguishes the reliable representations here. BIOT is a clear non-replication, reversing from +0.078 (95% CI +0.010 to +0.146) in Wang to -0.044 (-0.117 to +0.032) in Henao, and BENDR is consistently null at both intervals. BENDR’s final-stage collapse and the direction of LaBraM’s “an intermediate layer beats the final layer” pattern both replicate at Henao, though LaBraM’s specific peak layer does not, falling at block 1 in Henao versus block 8 in Wang, with overlapping confidence intervals. This directly motivated testing LaBraM’s entire layer-wise AUC curve on Miltiadous (Section 3.3) rather than committing to block 8 as a targeted claim.

**Figure 6.**
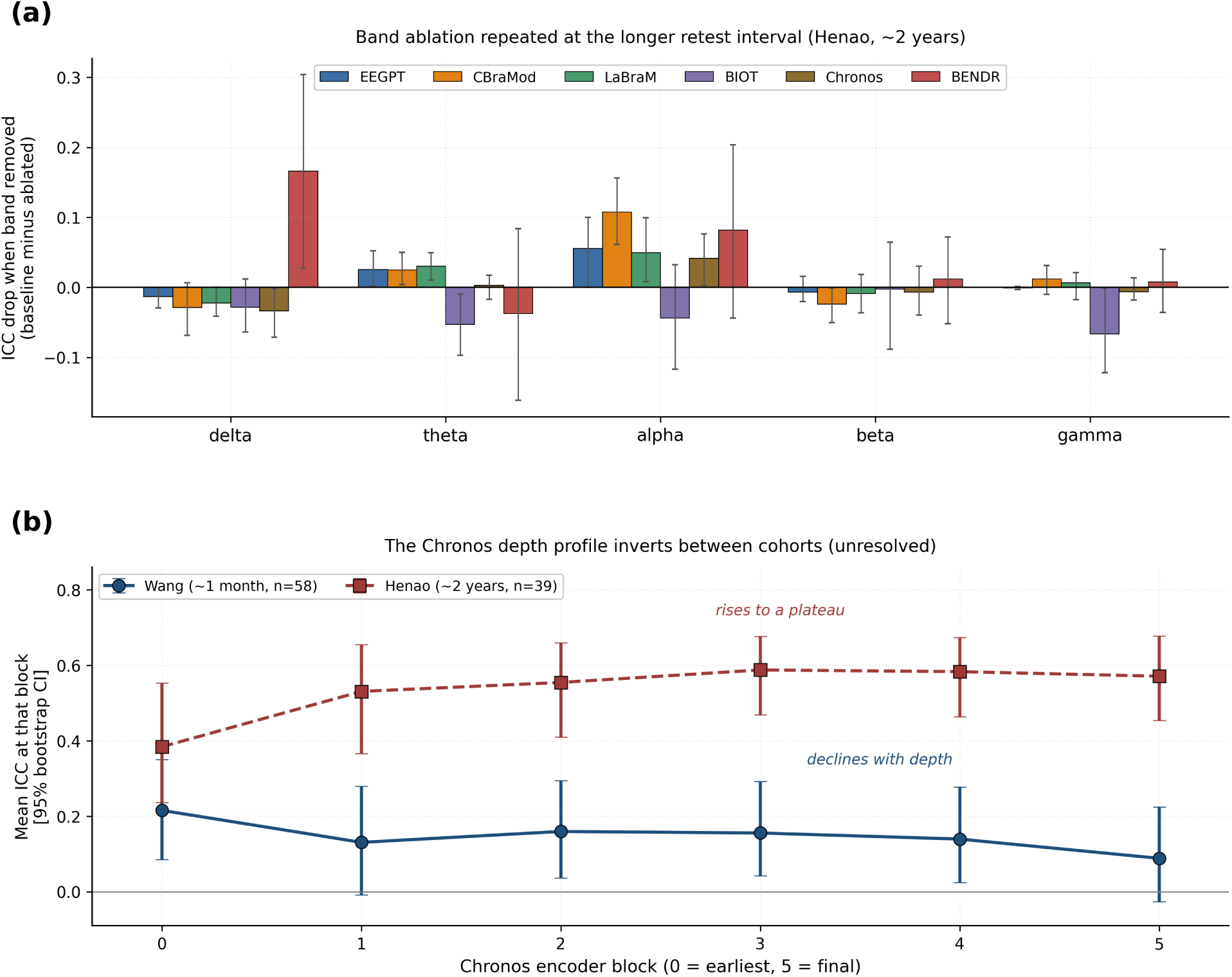
Cross-cohort replication at a longer retest interval, and the unresolved Chronos anomaly. (a) Frequency-band ablation repeated in the Henao cohort at an approximately two-year interval. Bars show the ICC drop when each band is removed, so positive values indicate that removing the band reduced reliability. The alpha effect replicates in CBraMod, EEGPT, LaBraM and Chronos, whose intervals exclude zero; BIOT reverses, and BENDR’s drop is positive but not significant at either interval. (b) Layer-wise reliability for Chronos in both cohorts, with 95% bootstrap confidence intervals. The entire depth profile inverts between cohorts: reliability is highest at the earliest encoder block and declines with depth in Wang, but is lowest at the earliest block and rises to a plateau in Henao, ending at 0.571 against 0.089 in Wang. Two candidate non-neural confounds, perrecording unit scaling and BIDS acquisition-metadata heterogeneity, were tested directly and ruled out (**Supplementary Table S16**). We report this as an open anomaly and do not treat Chronos’s Henao result as a replication.

Chronos shows a substantive, unresolved anomaly. Its baseline ICC increases from 0.075 in Wang, with a confidence interval crossing zero, to 0.571 (95% CI 0.444 to 0.678) in Henao, a 7.5-fold increase at the longer retest interval, the opposite direction from every other representation in this study. The entire layer-wise depth profile inverts between cohorts, not just the final-layer value (**Figure 6b**; per-block values in **Supplementary Table S15**). We directly tested and ruled out two candidate non-neural confounds: a per-recording unit-scaling artifact, and systematic between-subject differences in BIDS acquisition metadata. Neither can explain the anomaly (**Supplementary Table S16**). We report this prominently as a genuinely unresolved anomaly rather than attribute it to a specific confound, and we do not treat the Henao result as evidence that Chronos “also replicates.” This anomaly does not touch the paper’s other cross-cohort claims. Chronos appears throughout only as a non-EEG-pretraining comparison point, not as load-bearing evidence for the reliability-discrimination dissociation in Section 3.3, which rests on LaBraM, BENDR, and EEGPT for the geometric evidence and on all six perturbation-tested models jointly for the spectral evidence, nor for the architecture-depth typology in Section 3.4, where Chronos’s inverted profile is discussed separately rather than folded into the typology’s main claims.

## 4. Discussion

The two lines of evidence in Section 3.3 converge, and we think that convergence is the paper’s central contribution. A geometric analysis of embedding-space directions and a perturbational analysis of the input frequency spectrum, developed independently and using distinct methodology, both find that the information supporting test-retest reliability and the information supporting disease discrimination are not the same information. Neither analysis alone would be conclusive. The subspace-overlap result is correlational and limited to three models, and the band-ablation result rests on perturbations that change the input signal in several correlated ways. Together they are considerably more persuasive than either alone, because two structurally different methods failing to find shared information is harder to explain away as a shared methodological artifact than one method finding it once.

Synthesizing across all the evidence, this study supports a three-level account, summarised in **Figure 7**. At the spectral level, reliable representations preferentially encode alpha-band information that combines within-subject stability with between-subject distinctiveness, a candidate spectral contributor to reliability, while theta ranks first for disease discrimination instead, though only the reliability half of that contrast is individually significant. The “stability times individuality” pattern should be read as an alpha- and beta-specific observation, not a general principle. At the representational level, the dissociation is detectable both perturbationally and geometrically, by two independent routes. At the architectural level, whether this partial dissociation is preserved, traded off, or jointly degraded through network depth is itself architecture-specific: EEGPT and CBraMod preserve both properties, BENDR loses both at its final stage and does so at two retest intervals, and LaBraM shows a statistically supported reliability peak away from its final block, but no corresponding discrimination difference, so the tradeoff remains a point-estimate pattern rather than an established one.

**Figure 7.**
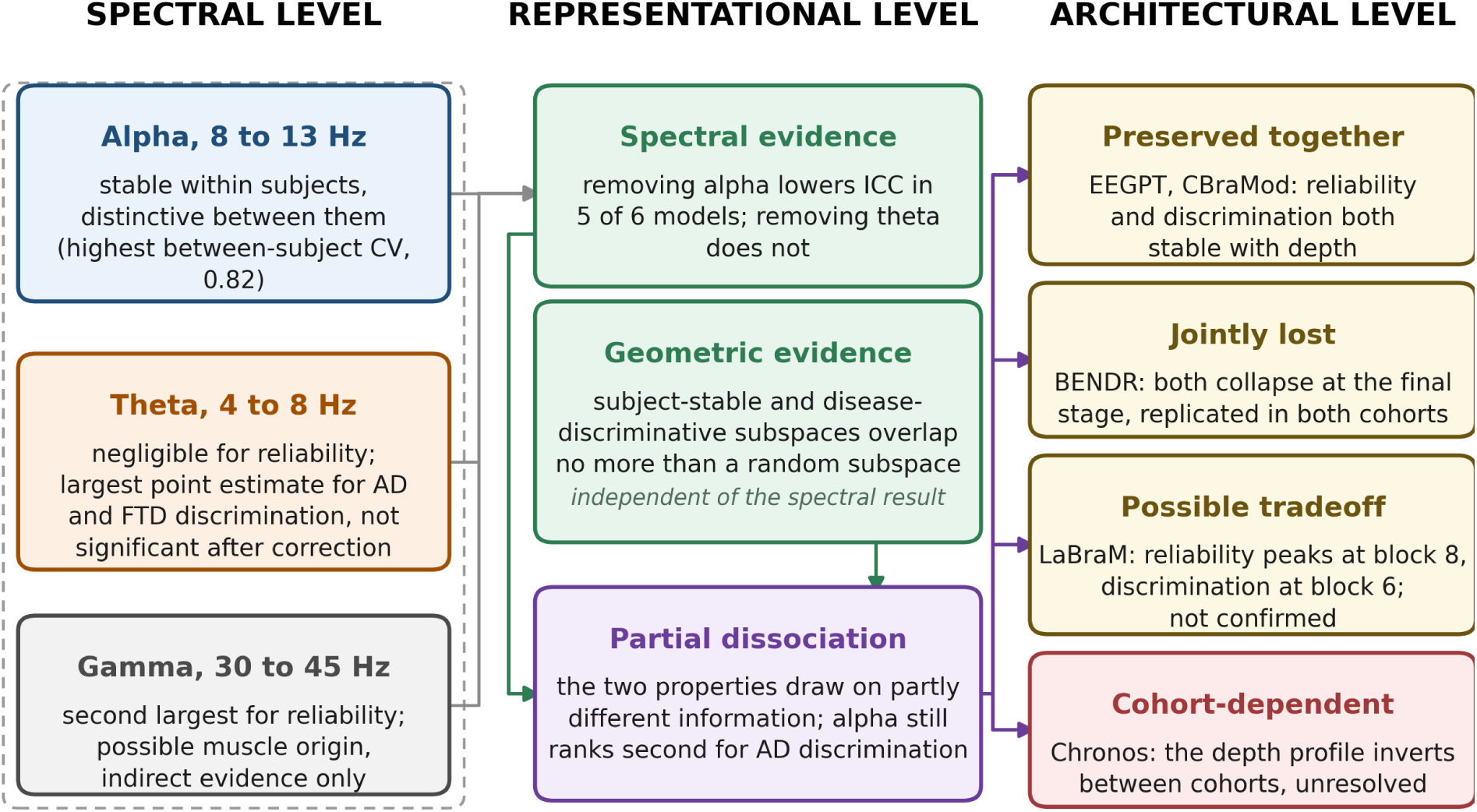
Conceptual overview: the three levels at which reliability and disease discrimination can be distinguished. Arrows indicate the flow of evidence, not causal influence. Left, spectral level: alpha-band information is both stable within subjects and distinctive between subjects, whereas theta-band information ranks first for disease discrimination by point estimate, though no individual band-by-model comparison survives multiplicity correction (**Supplementary Table S17**), and contributes negligibly to reliability; gamma is the second-largest reliability contributor, but its origin is uncertain and the evidence for it is indirect. Centre, representational level: two methodologically independent analyses converge on the same conclusion. Band ablation shows that removing alpha lowers reliability in five of six representations while removing theta does not, and a subspace analysis shows that the subject-stable and disease-discriminative directions of the embedding space overlap no more than a matched random-subspace null. Together these indicate a partial rather than a complete dissociation, since alpha still ranks second among bands for AD-versus-HC discrimination. Right, architectural level: whether an architecture preserves both properties across depth, loses both, or trades one against the other differs between models, and the Chronos depth profile inverts between cohorts and remains unresolved. Supporting values, with confidence intervals, are given in Sections 3.2 to 3.4.

The spread of reliability we observe, from an ICC near 0.08 up to 0.76, is not unusual by the standards of the wider neuroimaging reliability literature. Individual fMRI connectivity measurements span a comparably broad range, from near-zero to high, in a substantially larger evidence base (Noble et al., 2019), and the task-fMRI meta-analytic mean of 0.397 (Elliott et al., 2020) sits close to the middle of our own distribution. Wide variation in reliability appears to be a general property of multivariate neuroimaging measures, learned or handcrafted, rather than an idiosyncrasy of EEG foundation models specifically. The handcrafted end of our range is also externally calibrated: our alpha-power ICC of 0.867 at a one-month interval is close to the 0.843 reported for posterior alpha relative power across five years in an independent healthy cohort (Park et al., 2026), which suggests our pipeline is measuring the same underlying stability that the classical EEG literature measures, not an artifact of our particular preprocessing.

What is durable about our finding is structural, not a leaderboard: reliability varies widely and unpredictably across models that look equivalent on discrimination, and that structural claim should outlast any particular model’s ranking as architectures and training data continue to change.

The reliability paradox (Hedge et al., 2018) supplies a principled reason to expect the dissociation we measure, and our results extend it in a way that the original formulation does not anticipate. In the classical version, the tension is arithmetic: shrinking between-subject variance strengthens a group contrast and weakens an individual-differences measure, so the two trade off along a single axis. What we find is not a trade-off along one axis but a difference in content. No above-null overlap between the subject-stable and disease-discriminative subspaces was detected, and the frequency bands that carry each are different bands. A representation is not forced to choose between the two properties, as EEGPT and CBraMod demonstrate by holding both across depth. It is simply not guaranteed to preserve both, and nothing in a discrimination benchmark reveals which case a given model falls into.

Our findings also sit alongside a set of concurrent audits that reach compatible conclusions by other routes. Zare (2026b) reports that none of five EEG-FMs represented the long-range temporal correlations of the alpha envelope in temporal order, while all five were dominated by a recording-site axis. That study is careful about how far the result generalises: for the three raw-waveform models the probe recovered neither the temporal nor the spectral target and is therefore uninformative, so the spectral-temporal dissociation is established only for the two spectral-input models. Within that scope it is a dissociation between what these models represent well and what a transferable marker would require. The companion study (Zare, 2026a) uses band restriction as one of several targeted negative controls, and reports that dataset identity remains almost perfectly decodable from frozen embeddings even after band restriction and per-epoch normalisation. That is a different use of spectral manipulation from ours: it asks what a control cannot remove, whereas we ask which band a property depends on. We are not aware of prior work applying band ablation to the reliability axis. Lin et al. (2026) find subject-identity information pervasive enough in frozen representations to be mistaken for clinical signal. Our identifiability results qualify that picture usefully: identifiability and stability come apart, and LaBraM is the clearest case, highly distinctive (0.633) yet unstable (ICC 0.319). A model can encode who someone is without encoding a value that reproduces on retest, so a diagnostic that flags identity leakage does not by itself establish that a representation is usable longitudinally. That the most informative depth for downstream purposes need not be the final layer is already indicated for discrimination, through feature-encoding peaks at lower-to-middle depths (Tang et al., 2026) and through early blocks that already hold task-related information (Širca et al., 2026); what our layer-wise analysis adds is that the same is true for reliability, that the optimal depth for the two properties need not coincide, and that in one architecture reliability collapses at the final stage entirely. Širca et al. (2026) also show that poor head-only performance in EEG-FMs is largely attributable to pooling rather than to representational capacity. That result bears directly on ours, since it is consistent with our finding that BENDR’s and SignalJEPA’s distributed projection heads produce near-constant output while their convolutional encoders do not.

This has a direct practical implication. A representation or layer selected for reliability is not guaranteed to be optimal for disease discrimination, and applications that require both, such as a longitudinal disease biomarker that needs stability and disease sensitivity together, may need to combine information from multiple layers, or accept a compromise representation, rather than assume that a model’s default readout serves both purposes. It describes a property of the representations that a longitudinal biomarker application would still need to confirm in its intended patient population. There is also a more immediate, practical caution buried in Methods 2.3: for BENDR and Signal-JEPA, the distributed feature or projection head produced near-constant output across recordings, so a researcher loading either model out of the box would inherit a representation unusable for longitudinal tracking unless they manually route around the default head to the convolutional encoder, as we did here through a single uniform rule. More broadly, and echoing this study’s original negative finding, we found no detectable association between reliability and discrimination, pretraining paradigm, or pretraining domain. We present that negative result not as a standalone conclusion but as the setup for what we found in its place: a specific, falsifiable, partially generalising biological and architectural account of the factors that contribute to reliability, and of why they do not guarantee clinical usefulness on their own. The broader caution this motivates is that a learned embedding cannot be assumed suitable for longitudinal use simply because it comes from a foundation model, uses a particular pretraining objective, or classifies disease well. Woo et al. (2017) make the same argument for brain-based biomarkers generally: a candidate measure must be judged on reliability and generalisability, not on classification performance alone. Equivalence on a discrimination benchmark can conceal clinically important representational differences, a recognised hazard of benchmark-driven evaluation in medical machine learning (Varoquaux and Cheplygina, 2022), and two models that score alike on classification can still differ several-fold in whether they can track an individual over time.

Reliability is also distinct from representational distinctiveness (Section 3.1). A representation can encode who someone is without stably encoding their state, and only the latter supports tracking change over time, which is what a longitudinal biomarker requires. The fingerprinting literature establishes the first, that individuals are identifiable from short resting recordings (Demuru and Fraschini, 2020; da Silva Castanheira et al., 2021), without establishing the second. The reliability ordering of representations survived realistic acquisition degradation, which points to reliability as a property of the trained representation rather than a recording artifact, though robustness and baseline reliability are separable properties in their own right; BIOT, for instance, was perturbation-tolerant despite only intermediate baseline reliability.

Two directions follow most naturally from this work. The first is the obvious one: reliability must be measured in patients, not inferred from healthy cohorts. This is not a formality: reliability is a property of a measure in a population rather than of the measure alone (Koo and Li, 2016), and EEG reliability varies substantially with both (Lopez et al., 2023). The second is more tractable in the short term. If a subject-stable subspace can be recovered from LaBraM’s frozen features by a linear adapter, as we found, then reliability may be partly a property of the readout rather than of the pretrained weights, and architectures could be evaluated, or adapters trained, with reliability as an explicit objective rather than as an emergent accident of pretraining. The machinery exists but has been pointed the other way: adversarial objectives have been used to strip subject identity from EEG representations (Ozdenizci et al., 2020), and inverting them would preserve within-subject stability rather than suppress it. Neither direction requires new pretraining runs, which makes both accessible to groups without large compute budgets.

Two implications follow for how these models are built and assessed. For development, reliability is unusually cheap to report: it requires only repeated recordings and subject identifiers, with no disease labels and no clinical cohort, so it can be computed at pretraining time on any retest data rather than deferred to downstream clinical evaluation. A benchmark could require it alongside accuracy at essentially no additional annotation cost, and a developer could treat it as a target rather than discover it afterwards. For the evidentiary path toward clinical use, the FDA-NIH BEST framework separates analytical validation, whether a measurement is technically sound and reproducible, from clinical validation, whether it relates to the condition of interest (FDA-NIH Biomarker Working Group, 2016). Test-retest reliability is analytical-validation evidence, and what these results indicate is that it cannot be inferred from clinical-validation performance: the representations here are close to indistinguishable on the latter while differing several-fold on the former. We offer this as a correspondence between our measurements and an existing evidentiary vocabulary, not as a claim about what any particular qualification process would require.

## 5. Limitations

The limitations of this study fall into three groups: what the data cannot support, what the statistical treatment leaves open, and what the implementation choices constrain. We set them out in that order, beginning with the one that matters most.

Reliability was measured only in healthy adults, and this is the study’s central limitation. We examined two longitudinal ALS datasets to test patient generalisation, and both proved unsuitable. One yielded near-zero reliability for every representation, including handcrafted features, indicating that the recordings rather than the models were limiting. The other did not preserve the session structure that test-retest reliability requires. Openly downloadable disease test-retest EEG that is resting-state, repeated-session and adequately powered proved hard to obtain rather than nonexistent. Candidate resources do exist, including multi-site treatment trials that acquire resting EEG before and shortly after randomisation, and large protocol-driven cohorts with scheduled repeat recordings, but they variously sit behind data access applications, use low-density sleep montages, or confound the retest interval with a deliberate intervention. We did not pursue them here. Whether these patterns hold in patients is the primary direction for future work, and this gap is materially larger than any of the mechanistic limitations that follow. Sample sizes are also modest by neuroimaging-reliability standards, at Wang n = 58, Henao n = 39, and Miltiadous n = 85 or 63 for the AD-versus-HC contrast specifically, and several comparisons, including the layer-wise AUC paired tests and one PD cohort, are visibly underpowered.

An earlier planning stage considered a third masked-reconstruction model, LEAD, a window-length-matched comparability check, and a selection-criterion robustness analysis across multiple representation-selection rules. None of these three made it into the final evaluated set or the analyses reported here. Relatedly, the preprocessing-sensitivity comparison in Section 3.1 tests only one representation pair, handcrafted features against LaBraM, and does not test sensitivity to the window-length differences between handcrafted features and embeddings, which remains untested.

Multiple comparisons are addressed through predefined comparison families with within-family BH-FDR correction (Methods 2.4), not a single correction across the whole study. Two families with genuine permutation p-values, the AD-versus-HC and FTD-versus-HC band-ablation AUC tests, were corrected explicitly, and 18 of 18 and 29 of 36 tests respectively remain significant. The tests that do not survive correction concentrate in BENDR and do not touch the headline theta and alpha discrimination claims. The two layer-wise families are corrected within family, but their contents were specified after the layer-wise results were first inspected rather than in advance, so they are confirmatory only in the weak sense that the correction is applied to the full set of comparisons we report, not to every comparison the data would permit. Separately, nearly every representation-by-cohort combination was extracted only once. The one deliberate exception is the Henao BIOT layer-wise extraction, which was independently re-run from scratch and matched exactly; that is reassuring for the case it covers, but it is not evidence about extraction-procedure variance more broadly.

Why the models differ so much is largely beyond what these data can resolve, since pretraining objective, tokenisation, temporal context, and architecture are confounded across the models available today; isolating a causal design factor would require controlled training of matched architectures, which an evaluation study of this kind cannot provide.

The band-ablation results are perturbational evidence, not mechanistic proof. Band-stop filtering changes the input signal in multiple correlated ways, does not isolate a single biological mechanism, and introduces distortions that complicate interpretation of the filtered signal (de Cheveigné and Nelken, 2019). The supported claim is therefore the narrow one, that alpha-band information contributes disproportionately to representation reliability, and not that alpha-band neural activity causally determines individual identifiability.

Three implementation choices constrain interpretation. Layer-wise pooling is a generic heuristic, validated with a diagnostic pass but not independently verified against each model’s offiicial readout procedure. This limitation is not incidental, because Širca et al. (2026) show that pooling choice, rather than representational capacity, accounts for much of the apparent weakness of frozen EEG-FM readouts, so absolute layer-wise values here should be read as pooling-dependent and only the within-model depth trends as robust. For two models, BENDR and SignalJEPA, we used the convolutional-encoder representation rather than the distributed projection head (Methods 2.3), a principled but non-default choice. Model implementations and weights also continue to evolve, so we frame our results as findings about the spread and unpredictability of reliability across contemporary models rather than as a fixed leaderboard.

The spectral dissociation is asymmetric in how well its two halves are supported. Alpha’s contribution to reliability is backed by bootstrap intervals excluding zero in five of six representations, whereas theta’s contribution to discrimination reproduces as a ranking across both disease contrasts but yields no model-by-band comparison that survives Benjamini-Hochberg correction, and only eight of sixty intervals exclude zero uncorrected (**Supplementary Table S17**). The dissociation should therefore be read as well supported on the reliability axis and suggestive on the discrimination axis at this sample size.

MOMENT and Chronos were originally scored without the age and sex predictors supplied to the other seven representations, because their exported matrices lacked those columns. All nine now share one implementation (src/discrimination.py) and were rescored under it, permutation tests included. The harmonisation moved their AUCs by at most 0.022 and changed no significance verdict, but it did change one ranking: MOMENT is second rather than first of nine on the Cavanagh Parkinson’s contrast.

Two specific claims rest on weaker evidence than the rest and should be read accordingly. The gamma-artifact-contamination hypothesis is supported only indirectly, by two crude topographic and kurtosis proxies, and an ICA-based ground-truth decomposition would be needed for a stronger claim. The Chronos anomaly remains unresolved, and Chronos’s cross-cohort results should not be presented as a clean replication for any purpose until it is understood. Coverage is also uneven across models: the robustness stress test and the geometric mechanistic analyses of recoverability and subspace overlap were run only for LaBraM, BENDR, and EEGPT, and were not extended to MOMENT, Chronos, or the remaining EEG-oriented models, so whether MOMENT’s high reliability or Chronos’s anomalous values hold up under the same analyses is untested.

Finally, these analyses are exploratory throughout and were not pre-registered. They are hypothesis-generating, and they motivate a pre-registered confirmatory study, ideally in disease retest cohorts.

## 6. Conclusion

Across nine representations spanning the major EEG-FM pretraining paradigms and two general-purpose time-series models with no EEG exposure, test-retest reliability ranged from minimal to high, while disease discrimination remained comparable, and reliability’s variation was predicted by neither discrimination, pretraining paradigm, nor pretraining domain. We then identified a specific, partially generalising explanation for that variation. Across most representations tested, alpha-band information made a disproportionate contribution to reliability. This information is partially but not fully distinct from the information supporting disease discrimination, a conclusion supported by two independent methods. Whether this distinction is preserved, traded off, or jointly degraded depends on network architecture and depth. Measurement reliability is a separable, model-specific, and now partially characterised property, at the physiological, representational, and architectural levels, that must be established empirically for each model and representation layer, not assumed from category, pretraining domain, or classification accuracy. We recommend that test-retest reliability be reported as a standard evaluation axis alongside accuracy, and that representation or layer selection for longitudinal biomarker use account for the reliability-discrimination dissociation identified here. We release a reproducible pipeline for both the benchmarking and mechanistic analyses. Whether these patterns hold in patients, and what explains the still-unresolved Chronos anomaly, are the two most important open questions this study leaves for future work.

## Data and code availability

All analysis code, the unit-tested reliability implementation, the unified preprocessing pipeline, and a documented analysis protocol are at https://github.com/BelayTG/eeg-fm-reliability. Datasets are publicly available on OpenNeuro (ds004148, ds007176, ds004504, ds002778, ds003490).

## Ethical statement

This study used only pre-existing, fully de-identified, publicly available human EEG recordings distributed through OpenNeuro (ds004148, ds007176, ds004504, ds002778, ds003490). No new human-subjects data were collected, and no identifiable participant information was accessed at any point. Ethical approval and informed consent were obtained by the original investigators of each contributing cohort, as documented in the respective dataset records and accompanying data descriptors. Secondary analysis of these de-identified public datasets did not require additional ethical approval at the authors’ institutions. All analyses complied with the data-use terms attached to each OpenNeuro record.

## Acknowledgements

We thank the investigators who collected and openly shared the EEG datasets used here, and the developers of the foundation models and the braindecode, MNE-Python, and specparam software this work builds on.

## CRediT author contributions

**Belay Tadesse Gebregergis:** Conceptualization, Methodology, Software, Formal analysis, Investigation, Data curation, Writing – original draft, Writing – review and editing, Visualization. **Haben Girmay Yhdego:** Methodology, Software, Validation, Writing – review and editing. **Tewolde Teklu:** Conceptualization, Supervision, Resources, Writing – review and editing.

## Use of AI tools

Large language model assistants were used during the preparation of this manuscript for editing of the text, for debugging analysis code, and for literature search. All analyses were specified and run by the authors, who take full responsibility for the content.

