## Supplementary Material for "Reliability and disease sensitivity are dissociable properties of EEG foundation-model representations"

### Supplementary Tables

**Supplementary Table S1. Datasets and acquisition details.**
Five public OpenNeuro cohorts: two healthy test-retest cohorts (reliability analysis) and three neurodegenerative cohorts (discrimination analysis). Recording counts are post-quality-control. For every cohort the group column gives the published cohort size followed by the number that passed this study's unified quality-control pipeline, since the two differ slightly in all three cases where retest structure or exclusions apply.

| Cohort (OpenNeuro) | Role | Groups (n) | Sessions / interval | Channels | Sampling | Recordings analysed |
|---|---|---|---|---|---|---|
| Wang (ds004148) | Reliability | 60 healthy published; 59 passed QC, 58 with >=2 sessions | 3 sessions; two ~90 min apart, one ~1 month | 61 | 500 Hz | 167 |
| Henao Isaza (ds007176) | Reliability | 45 healthy published; 44 passed QC, 39 with >=2 sessions | up to 4 sessions over ~2 years | 60 | 1000 Hz | 141 |
| Miltiadous (ds004504) | Discrimination | 34 AD, 22 FTD, 29 HC post-QC (published dataset: 36 AD, 23 FTD, 29 HC) | single session | 19 | 500 Hz | 85 |
| Rockhill (ds002778) | Discrimination | ~15 PD, ~16 HC (OFF med) | single session | ~32 | 512 Hz | 31 |
| Cavanagh (ds003490) | Discrimination | 25 PD, 25 HC (OFF med) | single session | 63 | 500 Hz | 50 |

*Note: the Miltiadous row reports the post-quality-control analysed sample (34 AD, 22 FTD, 29 HC = 85 recordings), verified directly against `results/miltiadous_labels.csv`; the published dataset's full un-excluded cohort is 36 AD, 23 FTD, 29 HC. Three recordings (2 AD, 1 FTD) did not pass this study's unified QC pipeline (Methods 2.2) and are excluded from all analyses in this paper.*

**Supplementary Table S2. Frozen-embedding extraction and input-responsiveness validation.**
For each foundation model, the embedding extracted and a responsiveness check: the mean absolute change in the embedding across three very different random inputs, relative to its own standard deviation (a value near zero indicates a near input-invariant, unusable representation). For BENDR and SignalJEPA, the distributed models' downstream feature/projection heads were near input-invariant under frozen inference; we therefore applied a single uniform rule, use the convolutional feature-encoder representation, global-average-pooled over time, verified input-responsive for both.

| Model | Paradigm | Embedding used | Dim | Responsiveness ratio | Status |
|---|---|---|---|---|---|
| LaBraM | masked | mean-pooled tokens (final layer) | 200 | responsive | default API |
| CBraMod | masked | mean-pooled patch features | 200 | responsive | default API |
| EEGPT | contrastive | mean-pooled summary tokens | 2048 | responsive | default API |
| BIOT | masked/tokenized | default feature output (256-d), mean-pooled | 256 | responsive (cos-dist 0.10 across recordings) | default API |
| BENDR | contrastive | conv. encoder, GAP over time | 512 | 1.21 | encoder rule (default head near-constant across recordings; see Supplementary Table S3) |
| SignalJEPA | predictive | conv. encoder, GAP over time | 64 | 0.17 | encoder rule (default head constant; see Supplementary Table S3) |

**Supplementary Table S3. Why the default heads are unusable as frozen embeddings: head vs convolutional-encoder behaviour across real recordings.**

For BENDR and SignalJEPA we compare the model's default output head against the convolutional feature-encoder used in this study, on a sample of 18 real recordings (Wang cohort), plus a random-input responsiveness ratio. The decisive columns are the across-recording statistics: a representation useful for reliability must vary across recordings of different individuals. Both default heads collapse on real EEG; the convolutional encoders retain clear recording-to-recording structure.

| Representation | Random-input responsiveness ratio | Across-recording per-dim SD | Across-recording cosine distance: mean (SD) [min, max] |
|---|---|---|---|
| BENDR, default head | 0.25 | 1.13e-04 | 0.0000 (0.0000) [0.0000, 0.0000] |
| BENDR, conv. encoder (used) | 1.21 | 8.34e-03 | 0.0014 (0.0003) [0.0008, 0.0025] |
| SignalJEPA, default head | 0.00 (constant) | ~0 (constant) | constant output (undefined) |
| SignalJEPA, conv. encoder (used) | 0.17 | 3.16e-04 | 0.0049 (0.0038) [0.0013, 0.0256] |

**Supplementary Table S4. Pairwise reliability differences (paired bootstrap).**

Difference in mean ICC for sixteen predefined representation pairs, each computed on the subjects common to both representations using the same bootstrap resample (2000 iterations, independent seeded stream per pair); 95% CI on the difference. The pairs were chosen to cover each model against the handcrafted reference, agreement within a pretraining family, and the masked-versus-contrastive contrast across families; they are not the full set of 21 possible pairs. "Excludes 0" indicates a statistically supported difference.

*Wang (~1 month):*

| Pair | Delta mean ICC [95% CI] | n | Significance |
|---|---|---|---|
| Handcrafted - LaBraM | +0.431 [+0.331, +0.527] | 58 | excludes 0 |
| Handcrafted - CBraMod | +0.080 [+0.026, +0.129] | 58 | excludes 0 |

| Pair | Delta mean ICC [95% CI] | n | Significance |
|---|---|---|---|
| Handcrafted - EEGPT | -0.000 [-0.062, +0.055] | 58 | includes 0 (ns) |
| Handcrafted - BENDR | +0.618 [+0.491, +0.738] | 58 | excludes 0 |
| LaBraM - CBraMod | -0.353 [-0.440, -0.252] | 58 | excludes 0 |
| EEGPT - BENDR | +0.618 [+0.503, +0.728] | 58 | excludes 0 |
| EEGPT - LaBraM | +0.433 [+0.345, +0.521] | 58 | excludes 0 |
| EEGPT - CBraMod | +0.080 [+0.029, +0.134] | 58 | excludes 0 |
| BENDR - LaBraM | -0.188 [-0.335, -0.037] | 58 | excludes 0 |
| BENDR - CBraMod | -0.540 [-0.652, -0.430] | 58 | excludes 0 |
| Handcrafted - SignalJEPA | +0.334 [+0.245, +0.415] | 58 | excludes 0 |
| SignalJEPA - EEGPT | -0.336 [-0.410, -0.268] | 58 | excludes 0 |
| SignalJEPA - LaBraM | +0.097 [-0.007, +0.195] | 58 | includes 0 (ns) |
| Handcrafted - BIOT | +0.380 [+0.257, +0.497] | 58 | excludes 0 |
| BIOT - EEGPT | -0.380 [-0.486, -0.269] | 58 | excludes 0 |
| BIOT - CBraMod | -0.297 [-0.406, -0.187] | 58 | excludes 0 |

*Henao Isaza (~2 years):*

| Pair | Delta mean ICC [95% CI] | n | Significance |
|---|---|---|---|
| Handcrafted - LaBraM | +0.166 [+0.049, +0.261] | 34 | excludes 0 |
| Handcrafted - CBraMod | +0.024 [-0.081, +0.109] | 34 | includes 0 (ns) |
| Handcrafted - EEGPT | +0.058 [-0.138, +0.196] | 34 | includes 0 (ns) |
| Handcrafted - BENDR | +0.473 [+0.304, +0.609] | 34 | excludes 0 |
| LaBraM - CBraMod | -0.122 [-0.188, -0.047] | 39 | excludes 0 |
| EEGPT - BENDR | +0.462 [+0.307, +0.620] | 39 | excludes 0 |
| EEGPT - LaBraM | +0.078 [+0.010, +0.162] | 39 | excludes 0 |
| EEGPT - CBraMod | -0.042 [-0.115, +0.044] | 39 | includes 0 (ns) |
| BENDR - LaBraM | -0.378 [-0.515, -0.239] | 39 | excludes 0 |
| BENDR - CBraMod | -0.504 [-0.630, -0.374] | 39 | excludes 0 |
| Handcrafted - SignalJEPA | +0.334 [+0.175, +0.449] | 34 | excludes 0 |
| SignalJEPA - EEGPT | -0.270 [-0.333, -0.207] | 39 | excludes 0 |
| SignalJEPA - LaBraM | -0.189 [-0.262, -0.102] | 39 | excludes 0 |
| Handcrafted - BIOT | +0.133 [+0.032, +0.236] | 34 | excludes 0 |
| BIOT - EEGPT | -0.067 [-0.217, +0.058] | 39 | includes 0 (ns) |
| BIOT - CBraMod | -0.109 [-0.206, -0.015] | 39 | excludes 0 |

**Supplementary Table S5. Full disease-discrimination results.**
Permutation-validated repeated stratified 5-fold cross-validation AUC for each representation and contrast, with permutation-null mean and p-value (200 permutations). Benjamini-Hochberg FDR correction (q<0.05) across all 36 rows in this table did not change which results are significant (27/36 significant before and after correction; Methods 2.4).

| Cohort | Contrast | Representation | AUC | Null mean | Perm. p |
|---|---|---|---|---|---|
| Miltiadous | AD vs HC | Handcrafted | 0.849 | 0.505 | 0.005 |
| Miltiadous | AD vs HC | EEGPT | 0.876 | 0.510 | 0.005 |
| Miltiadous | AD vs HC | CBraMod | 0.854 | 0.492 | 0.005 |
| Miltiadous | AD vs HC | SignalJEPA | 0.884 | 0.497 | 0.005 |
| Miltiadous | AD vs HC | LaBraM | 0.899 | 0.504 | 0.005 |
| Miltiadous | AD vs HC | BENDR | 0.790 | 0.503 | 0.005 |
| Miltiadous | AD vs HC | BIOT | 0.858 | 0.497 | 0.005 |
| Miltiadous | AD vs HC | MOMENT | 0.801 | 0.495 | 0.005 |
| Miltiadous | AD vs HC | Chronos | 0.859 | 0.504 | 0.005 |
| Miltiadous | FTD vs HC | Handcrafted | 0.834 | 0.505 | 0.005 |
| Miltiadous | FTD vs HC | EEGPT | 0.770 | 0.497 | 0.020 |
| Miltiadous | FTD vs HC | CBraMod | 0.778 | 0.503 | 0.010 |

| Cohort | Contrast | Representation | AUC | Null mean | Perm. p |
|---|---|---|---|---|---|
| Miltiadous | FTD vs HC | SignalJEPA | 0.790 | 0.504 | 0.005 |
| Miltiadous | FTD vs HC | LaBraM | 0.792 | 0.501 | 0.005 |
| Miltiadous | FTD vs HC | BENDR | 0.694 | 0.518 | 0.035 |
| Miltiadous | FTD vs HC | BIOT | 0.798 | 0.500 | 0.005 |
| Miltiadous | FTD vs HC | MOMENT | 0.763 | 0.502 | 0.010 |
| Miltiadous | FTD vs HC | Chronos | 0.773 | 0.509 | 0.005 |
| Cavanagh | PD vs HC | Handcrafted | 0.795 | 0.493 | 0.005 |
| Cavanagh | PD vs HC | EEGPT | 0.803 | 0.487 | 0.005 |
| Cavanagh | PD vs HC | CBraMod | 0.724 | 0.501 | 0.005 |
| Cavanagh | PD vs HC | SignalJEPA | 0.380 | 0.488 | 0.856 |
| Cavanagh | PD vs HC | LaBraM | 0.696 | 0.500 | 0.025 |
| Cavanagh | PD vs HC | BENDR | 0.791 | 0.485 | 0.005 |
| Cavanagh | PD vs HC | BIOT | 0.826 | 0.497 | 0.010 |
| Cavanagh | PD vs HC | MOMENT | 0.821 | 0.479 | 0.005 |
| Cavanagh | PD vs HC | Chronos | 0.809 | 0.493 | 0.010 |
| Rockhill | PD vs HC | Handcrafted | 0.543 | 0.500 | 0.388 |
| Rockhill | PD vs HC | EEGPT | 0.346 | 0.492 | 0.851 |
| Rockhill | PD vs HC | CBraMod | 0.351 | 0.484 | 0.856 |
| Rockhill | PD vs HC | SignalJEPA | 0.424 | 0.487 | 0.627 |
| Rockhill | PD vs HC | LaBraM | 0.393 | 0.497 | 0.751 |
| Rockhill | PD vs HC | BENDR | 0.783 | 0.502 | 0.025 |
| Rockhill | PD vs HC | BIOT | 0.372 | 0.502 | 0.836 |
| Rockhill | PD vs HC | MOMENT | 0.522 | 0.504 | 0.453 |
| Rockhill | PD vs HC | Chronos | 0.489 | 0.510 | 0.582 |

**Supplementary Table S6. Full test-retest reliability with bootstrap confidence intervals (both cohorts).** Mean ICC(2,1) over representation dimensions, longest-interval pair per subject; 95% CIs from 2000 subject-level bootstrap iterations (seed 42).

| Representation (paradigm) | Wang ICC [95% CI] (n=58) | Henao ICC [95% CI] (n=34-39) |
|---|---|---|
| Handcrafted | 0.760 [0.668, 0.820] | 0.653 [0.459, 0.734] |
| EEGPT (contrastive) | 0.756 [0.678, 0.805] | 0.609 [0.481, 0.744] |
| CBraMod (masked) | 0.676 [0.596, 0.733] | 0.663 [0.534, 0.752] |
| MOMENT (general time series) | 0.586 [0.487, 0.669] | 0.696 [0.566, 0.766] |
| SignalJEPA (predictive) | 0.420 [0.333, 0.480] | 0.334 [0.218, 0.467] |
| BIOT (masked/tokenized) | 0.373 [0.268, 0.473] | 0.555 [0.433, 0.641] |
| LaBraM (masked) | 0.319 [0.228, 0.407] | 0.533 [0.429, 0.635] |
| BENDR (contrastive) | 0.133 [0.032, 0.221] | 0.152 [0.044, 0.239] |
| Chronos (general time series) | 0.075 [-0.038, 0.212] | 0.571 [0.448, 0.680] |

**Supplementary Table S7. Multivariate (dimensionality-agnostic) reliability and identifiability, both cohorts.** Within/between representational distance ratios, lower meaning more reliable, with 95% jackknife CIs, ordered by mean ICC. The agreement ratio uses Euclidean distance on cohort-standardised dimensions and is the multivariate counterpart of the absolute-agreement ICC(2,1); the consistency ratio uses correlation distance, which is invariant to shifting or rescaling a whole recording (Methods 2.4). The agreement ratio tracks the ICC ordering in both cohorts (Spearman rho = -0.85, p = 0.004 in Wang; rho = -0.75, p = 0.020 in Henao Isaza). The consistency ratio tracks it in Wang (rho = -0.90, p = 0.001) but not in Henao Isaza (rho = -0.45, p = 0.224), where CBraMod ranks second by ICC and eighth of nine by that ratio, which is why the consistency variant is not used to corroborate an absolute-agreement measure. Identifiability is the fraction of recordings whose nearest neighbour is another session of the same subject, chance 0.011 in Wang; it is a distinctiveness measure, not a reliability measure.

*Wang (~1 month):*

| Representation | Dim | Mean ICC | Agreement ratio [95% CI] | Consistency ratio [95% CI] | Identifiability [95% CI] |
|---|---|---|---|---|---|
| Handcrafted | 13 | 0.760 | 0.450 [0.388, 0.513] | 0.174 [0.101, 0.246] | 0.229 [0.118, 0.340] |
| EEGPT | 2048 | 0.756 | 0.460 [0.404, 0.515] | 0.233 [0.144, 0.321] | 0.711 [0.609, 0.813] |
| CBraMod | 200 | 0.676 | 0.544 [0.492, 0.596] | 0.481 [0.284, 0.678] | 0.602 [0.492, 0.713] |
| MOMENT | 1024 | 0.586 | 0.525 [0.462, 0.589] | 0.306 [0.228, 0.385] | 0.500 [0.396, 0.604] |
| SignalJEPA | 64 | 0.420 | 0.765 [0.724, 0.807] | 0.506 [0.431, 0.580] | 0.120 [0.042, 0.199] |
| BIOT | 256 | 0.373 | 0.603 [0.553, 0.653] | 0.472 [0.382, 0.561] | 0.572 [0.480, 0.664] |
| LaBraM | 200 | 0.319 | 0.639 [0.587, 0.691] | 0.556 [0.473, 0.640] | 0.633 [0.540, 0.725] |
| BENDR | 512 | 0.133 | 0.946 [0.914, 0.977] | 0.905 [0.876, 0.934] | 0.018 [0.000, 0.064] |
| Chronos | 512 | 0.075 | 0.683 [0.618, 0.749] | 0.703 [0.584, 0.823] | 0.422 [0.310, 0.534] |

*Henao Isaza (~2 years):*

| Representation | Dim | Mean ICC | Agreement ratio [95% CI] | Consistency ratio [95% CI] | Identifiability [95% CI] |
|---|---|---|---|---|---|
| MOMENT | 1024 | 0.696 | 0.499 [0.424, 0.574] | 0.220 [0.133, 0.307] | 0.397 [0.264, 0.530] |
| CBraMod | 200 | 0.663 | 0.719 [0.626, 0.812] | 0.872 [0.701, 1.043] | 0.412 [0.293, 0.530] |
| Handcrafted | 13 | 0.653 | 0.563 [0.484, 0.641] | 0.390 [-0.048, 0.828] | 0.213 [0.110, 0.317] |
| EEGPT | 2048 | 0.609 | 0.711 [0.628, 0.793] | 0.762 [0.596, 0.927] | 0.610 [0.507, 0.713] |
| Chronos | 512 | 0.571 | 0.558 [0.492, 0.624] | 0.278 [0.191, 0.365] | 0.346 [0.226, 0.465] |
| BIOT | 256 | 0.555 | 0.691 [0.624, 0.759] | 0.493 [0.363, 0.624] | 0.419 [0.282, 0.557] |
| LaBraM | 200 | 0.533 | 0.759 [0.675, 0.844] | 0.709 [0.558, 0.861] | 0.485 [0.353, 0.617] |
| SignalJEPA | 64 | 0.334 | 0.861 [0.807, 0.914] | 0.721 [0.617, 0.825] | 0.103 [0.049, 0.157] |
| BENDR | 512 | 0.152 | 0.944 [0.909, 0.979] | 0.968 [0.935, 1.001] | 0.037 [0.000, 0.087] |

**Supplementary Table S8. Matched-dimensionality control (PCA).**

Mean ICC computed over the leading k principal components (PCA fit on pooled two-session data), for each representation at matched dimensionalities. Entries marked n/a are cases where k exceeds the representation's native dimensionality, which is 13 for handcrafted features and 64 for SignalJEPA. The number of components a PCA can return is also bounded by the number of samples, 116 in Wang and 78 in Henao, so in Henao the k = 100 and k = 200 rows are both truncated to the same maximum and are identical by construction rather than by coincidence.

*Wang:*

| k | Handcrafted | LaBraM | CBraMod | EEGPT | BENDR | SignalJEPA | BIOT | MOMENT | Chronos |
|---|---|---|---|---|---|---|---|---|---|
| 5 | 0.723 | 0.515 | 0.670 | 0.747 | 0.236 | 0.334 | 0.360 | 0.545 | 0.328 |

| k | Handcrafted | LaBraM | CBraMod | EEGPT | BENDR | SignalJEPA | BIOT | MOMENT | Chronos |
|---|---|---|---|---|---|---|---|---|---|
| 10 | 0.651 | 0.476 | 0.561 | 0.709 | 0.201 | 0.273 | 0.372 | 0.576 | 0.286 |
| 13 | 0.617 | 0.458 | 0.492 | 0.676 | 0.177 | 0.254 | 0.339 | 0.532 | 0.300 |
| 50 | n/a | 0.251 | 0.220 | 0.447 | 0.039 | 0.082 | 0.250 | 0.294 | 0.180 |
| 100 | n/a | 0.050 | 0.036 | 0.093 | 0.012 | n/a | 0.072 | 0.055 | 0.035 |
| 200 | n/a | -0.002 | 0.002 | 0.002 | 0.001 | n/a | -0.002 | 0.005 | 0.001 |

*Henao:*

| k | Handcrafted | LaBraM | CBraMod | EEGPT | BENDR | SignalJEPA | BIOT | MOMENT | Chronos |
|---|---|---|---|---|---|---|---|---|---|
| 5 | 0.668 | 0.537 | 0.550 | 0.609 | 0.222 | 0.378 | 0.520 | 0.594 | 0.501 |
| 10 | 0.471 | 0.513 | 0.510 | 0.531 | 0.168 | 0.200 | 0.442 | 0.553 | 0.441 |
| 13 | 0.419 | 0.526 | 0.450 | 0.516 | 0.099 | 0.139 | 0.405 | 0.519 | 0.390 |
| 50 | n/a | 0.138 | 0.107 | 0.288 | 0.035 | 0.032 | 0.170 | 0.153 | 0.121 |
| 100 | n/a | 0.002 | 0.005 | -0.001 | 0.003 | n/a | 0.004 | 0.006 | 0.004 |
| 200 | n/a | 0.002 | 0.005 | -0.001 | 0.003 | n/a | 0.004 | 0.006 | 0.004 |

**Supplementary Table S9. Reliability stress test under simulated acquisition degradation.**
Mean per-dimension ICC across the Wang cohort, on the 54 subjects for whom every condition could be regenerated, for the six EEG-oriented models and handcrafted features under baseline and six perturbations. The final column, headed rank corr., is the Spearman rank correlation between each condition's ordering of the representations and the baseline ordering, so a value near 1 means the perturbation changed the absolute values without reordering the representations.

| Condition | Handcrafted | EEGPT | CBraMod | SignalJEPA | BIOT | LaBraM | BENDR | Rank corr. |
|---|---|---|---|---|---|---|---|---|
| baseline | 0.763 | 0.756 | 0.676 | 0.431 | 0.339 | 0.308 | 0.151 | reference |
| electrode loss 25% | 0.759 | 0.748 | 0.681 | 0.427 | 0.331 | 0.310 | 0.150 | 1.000 |
| electrode loss 50% | 0.746 | 0.736 | 0.688 | 0.423 | 0.292 | 0.321 | 0.175 | 0.964 |
| duration 50% | 0.773 | 0.750 | 0.640 | 0.368 | 0.371 | 0.329 | 0.129 | 0.964 |
| duration 25% | 0.757 | 0.725 | 0.554 | 0.293 | 0.364 | 0.322 | 0.020 | 0.893 |
| noise 0.5x | 0.767 | 0.759 | 0.701 | 0.422 | 0.432 | 0.619 | 0.208 | 0.857 |
| noise 1.0x | 0.731 | 0.764 | 0.750 | 0.391 | 0.474 | 0.632 | 0.104 | 0.750 |

**Supplementary Table S10. Variance-component decomposition under heavy noise.**
Used to diagnose whether the apparent ICC increases for LaBraM and CBraMod under added noise reflect a real reliability improvement or variance compression, on the same 54 Wang subjects, with noise at 1.0 times the per-channel standard deviation. Within and between denote the within-subject and between-subject standard deviations; base is the unperturbed condition and noise the perturbed one, and the two percentage columns give the change from base to noise. Compression, meaning both standard deviations falling together, raises ICC without any genuine gain in reliability. Reading across the representations: handcrafted features and EEGPT are stable, with neither standard deviation moving appreciably. LaBraM shows strong compression, the within-subject and between-subject values falling by 82% and 72% respectively, which is what produces its apparent ICC gain. BIOT and SignalJEPA show mild compression. CBraMod's between-subject standard deviation rises rather than falls, so its ICC change is not compression. BENDR is unaffected because it is already collapsed at baseline.

| Representation | Within (base) | Between (base) | Within (noise) | Between (noise) | Within % | Between % |
|---|---|---|---|---|---|---|
| Handcrafted | 0.0292 | 0.1211 | 0.0305 | 0.1307 | +5 | +8 |
| EEGPT | 0.0546 | 0.2065 | 0.0510 | 0.1972 | -7 | -5 |
| CBraMod | 0.0004 | 0.0013 | 0.0005 | 0.0018 | +7 | +33 |
| LaBraM | 0.2656 | 0.4669 | 0.0473 | 0.1305 | -82 | -72 |

| Representation | Within (base) | Between (base) | Within (noise) | Between (noise) | Within % | Between % |
|---|---|---|---|---|---|---|
| BIOT | 0.3538 | 0.6947 | 0.2903 | 0.6406 | -18 | -8 |
| SignalJEPA | 0.0001 | 0.0003 | 0.0001 | 0.0003 | -7 | -12 |
| BENDR | 0.0042 | 0.0063 | 0.0042 | 0.0060 | 0 | -6 |

**Supplementary Table S11. Preprocessing-sensitivity comparison (Wang).**
Mean ICC under the full unified pipeline versus minimal preprocessing (band-pass, average reference, unit normalisation only), for handcrafted features and LaBraM, the only embedding model tested under this comparison. Referenced in Section 3.1 of the main manuscript.

| Representation | Minimal preprocessing | Full pipeline |
|---|---|---|
| Handcrafted | 0.522 | 0.760 |
| LaBraM | 0.449 | 0.319 |

**Supplementary Table S12. Full band-by-band reliability-ablation breakdown (Wang cohort, 6 representatives).**
ICC drop (baseline minus band-ablated) with 95% bootstrap CI (n_boot=1000, seed 42), for every band tested, not just alpha. Several non-alpha effects reach significance here. Significance in this table means only that a confidence interval excludes zero; it is not evidence that a band is a major determinant of reliability. The case for alpha rests on effect magnitude, on consistency across representations, and on the ordering being reproduced in a second cohort, none of which the smaller significant effects satisfy. Referenced from Section 3.2 of the main manuscript, which reports only the alpha row in the main text; this table gives the full picture, including the two cases (BIOT, Chronos) where gamma's point estimate marginally exceeds alpha's.

| Representation | Band removed | ICC drop | 95% CI | Significant (CI excludes 0)? |
|---|---|---|---|---|
| BENDR | alpha | 0.058 | [-0.045, 0.150] | no |
| BENDR | beta | -0.039 | [-0.103, 0.021] | no |
| BENDR | delta | 0.062 | [-0.074, 0.197] | no |
| BENDR | gamma | 0.024 | [-0.022, 0.066] | no |
| BENDR | theta | 0.003 | [-0.081, 0.089] | no |
| BIOT | alpha | 0.078 | [0.010, 0.146] | yes |
| BIOT | beta | -0.015 | [-0.076, 0.050] | no |
| BIOT | delta | 0.005 | [-0.018, 0.027] | no |
| BIOT | gamma | 0.128 | [0.061, 0.184] | yes |
| BIOT | theta | -0.013 | [-0.053, 0.030] | no |
| CBraMod | alpha | 0.094 | [0.035, 0.156] | yes |
| CBraMod | beta | -0.012 | [-0.039, 0.008] | no |
| CBraMod | delta | -0.062 | [-0.099, -0.026] | yes (negative: removing delta *raised* ICC) |
| CBraMod | gamma | 0.007 | [-0.011, 0.025] | no |
| CBraMod | theta | 0.009 | [0.0002, 0.020] | yes (small effect) |
| Chronos | alpha | 0.058 | [0.025, 0.095] | yes |
| Chronos | beta | -0.013 | [-0.033, 0.009] | no |
| Chronos | delta | -0.023 | [-0.039, -0.007] | yes (negative) |
| Chronos | gamma | 0.083 | [0.054, 0.105] | yes |
| Chronos | theta | -0.013 | [-0.019, -0.005] | yes (negative) |
| EEGPT | alpha | 0.084 | [0.035, 0.134] | yes |
| EEGPT | beta | 0.016 | [0.001, 0.030] | yes (small effect) |
| EEGPT | delta | 0.000 | [-0.007, 0.006] | no |
| EEGPT | gamma | -0.003 | [-0.005, -0.001] | yes (small negative) |
| EEGPT | theta | 0.005 | [-0.002, 0.013] | no |

| Representation | Band removed | ICC drop | 95% CI | Significant (CI excludes 0)? |
|---|---|---|---|---|
| LaBraM | alpha | 0.148 | [0.103, 0.191] | yes |
| LaBraM | beta | 0.006 | [-0.016, 0.026] | no |
| LaBraM | delta | -0.006 | [-0.023, 0.010] | no |
| LaBraM | gamma | 0.023 | [0.006, 0.038] | yes (small effect) |
| LaBraM | theta | 0.024 | [0.007, 0.042] | yes (small effect) |

Note several statistically "significant" non-alpha effects here are small in magnitude (e.g. CBraMod/theta, EEGPT/beta, EEGPT/gamma) and are not treated as evidence of a competing mechanism in the main text; alpha and, secondarily, gamma remain the two bands with consistently large point estimates across representations.

**Supplementary Table S13. Subspace independence: principal-angle overlap between the subject-stable, within-subject noise, and disease-discriminative subspaces (0=orthogonal, 1=identical).**

The subject-stable and within-subject noise subspaces are the top five between-subject and within-subject principal directions in Wang. The disease-discriminative subspace is the between-class subspace of the Miltiadous class means, which for three classes has rank two; it is built at that rank rather than padded to five, since padding produced directions that were numerically degenerate rather than disease-related (Methods 2.4). Random null is the 95th percentile of the overlap between a random subspace of the same dimension as the stable subspace and the same rank-two disease subspace, over 200 draws, so the observed and null comparisons average over the same number of principal angles.

| Representation | Stable vs disease | Noise vs disease | Random null (95th pct) | Stable above null? |
|---|---|---|---|---|
| LaBraM | 0.187 | 0.166 | 0.191 | no (within null) |
| BENDR | 0.089 | 0.097 | 0.126 | no (within null) |
| EEGPT | 0.053 | 0.048 | 0.063 | no (within null) |

**Supplementary Table S14. Reliability recoverability: per-fold held-out test-retest ICC before and after a subject-invariance adapter.**

Five-fold subject-level cross-validation, seed 42. Every cell reads as baseline value then adapted value, and the change column is the difference in the fold mean.

| Representation | Fold 1 | Fold 2 | Fold 3 | Fold 4 | Fold 5 | Mean | Change |
|---|---|---|---|---|---|---|---|
| LaBraM | 0.121->0.570 | 0.484->0.496 | 0.252->0.478 | 0.296->0.418 | 0.308->0.468 | 0.292->0.486 | +0.194 |
| BENDR | 0.027->-0.011 | 0.252->0.170 | 0.145->0.066 | -0.016->0.061 | 0.204->0.092 | 0.122->0.076 | -0.047 |
| EEGPT | 0.763->0.688 | 0.806->0.706 | 0.670->0.672 | 0.636->0.671 | 0.773->0.656 | 0.730->0.679 | -0.051 |

**Supplementary Table S15. Chronos layer-wise reliability by encoder block, both cohorts.**

Mean per-dimension ICC with 95% bootstrap confidence intervals at each Chronos encoder block, extracted by the same forward-hook procedure used for the other layer-wise analyses. Block 5 is the default final layer whose values appear in Table 1. The two cohorts do not merely differ in magnitude: in Wang reliability is highest at the earliest block and declines with depth, whereas in Henao the earliest block is the least reliable and reliability rises to a plateau. The whole depth profile inverts, which argues against an explanation localised to the pooling convention at any single layer.

| Encoder block | Wang ICC [95% CI] | Henao ICC [95% CI] |
|---|---|---|
| 0 | 0.216 [0.085, 0.351] | 0.384 [0.237, 0.553] |
| 1 | 0.131 [-0.008, 0.279] | 0.531 [0.366, 0.655] |

| Encoder block | Wang ICC [95% CI] | Henao ICC [95% CI] |
|---|---|---|
| 2 | 0.160 [0.037, 0.295] | 0.555 [0.410, 0.660] |
| 3 | 0.156 [0.043, 0.293] | 0.588 [0.469, 0.677] |
| 4 | 0.140 [0.025, 0.278] | 0.583 [0.463, 0.674] |
| 5 (final, default) | 0.089 [-0.026, 0.225] | 0.571 [0.454, 0.678] |

**Supplementary Table S16. Non-neural confounds tested against the Chronos cross-cohort anomaly, and their outcomes.**

Chronos's baseline ICC rises from 0.075 in Wang, at roughly one month with a confidence interval crossing zero, to 0.571 in Henao at roughly two years, which is the opposite direction to every other representation in this study. Two candidate non-neural explanations were tested directly against the raw data rather than argued away. The BIDS check read the Henao ds007176 sidecar files covering 150 recordings from 45 subjects, 39 of whom have two or more sessions, and asked of each field whether it was consistent within a subject but variable between subjects, the pattern that could inflate ICC for a metadata-sensitive model. Full output is released as `tier2_chronos_anomaly_bids_metadata.csv`. Both confounds are excluded; no positive explanation has been identified, and the anomaly remains open.

| Candidate confound | What was tested | Outcome |
|---|---|---|
| Per-recording unit scaling | Whether the per-recording `unit_factor` term in the Chronos tokenisation pipeline varies between subjects rather than being fixed by recording hardware | Effectively constant within each subject in both cohorts, leaving no between-subject variance available to inflate ICC. Excluded. |
| Acquisition metadata: Manufacturer, EEGReference, PowerLineFrequency, SamplingFrequency, EEGGround | Whether any field is within-subject consistent but between-subject variable | Each is constant across the whole dataset, a single distinct value, so there is no between-subject heterogeneity at all. Excluded. |
| Acquisition metadata: RecordingDuration | Same test | Varies across recordings, but not consistently per subject, so it is not a stable between-subject signal a model could lock onto. Excluded. |

**Supplementary Table S17. Paired bootstrap on the band-ablation AUC drop.**

The permutation test reported in Supplementary Table S5 and in Table 2 shuffles diagnostic labels, so it asks whether an ablated representation still discriminates above chance. It does not ask whether removing a band changed the AUC. This table supplies that missing test, using the same design as the reliability side: out-of-fold predicted probabilities from the repeated stratified 5-fold pipeline, then 2000 subject-level resamples applied identically to the baseline and ablated conditions, stratified by class, with a two-sided percentile p and Benjamini-Hochberg correction within each contrast (30 tests per contrast, six representations by five bands). Baseline AUCs differ in the third decimal from Table 2 because the AUC is computed once on pooled out-of-fold predictions rather than averaged across fold-wise AUCs. Theta ranks first for both contrasts, matching the point-estimate ranking in Table 2, and accounts for five of the eight individually significant drops, but no test survives family-wise correction.

*AD versus HC:*

| Band removed | Mean AUC drop | Models with CI excluding 0 | Smallest p | Smallest p after BH |
|---|---|---|---|---|
| theta | +0.061 | 3 of 6 | 0.005 | 0.150 |
| alpha | +0.036 | 0 of 6 | 0.107 | 0.401 |
| beta | +0.016 | 0 of 6 | 0.084 | 0.401 |
| gamma | +0.011 | 0 of 6 | 0.125 | 0.416 |
| delta | +0.001 | 0 of 6 | 0.059 | 0.401 |

*FTD versus HC:*

| Band removed | Mean AUC drop | Models with CI excluding 0 | Smallest p | Smallest p after BH |
|---|---|---|---|---|
| theta | +0.061 | 2 of 6 | 0.030 | 0.192 |
| beta | +0.030 | 1 of 6 | 0.026 | 0.192 |
| gamma | +0.017 | 0 of 6 | 0.108 | 0.360 |
| alpha | +0.004 | 2 of 6 | 0.004 | 0.120 |
| delta | -0.024 | 0 of 6 | 0.068 | 0.291 |

**Supplementary Table S18. Age and sex alone, as a reference for the discrimination AUCs.**
Age and sex are appended to the embedding as additional predictors rather than partialled out, so they can contribute to the reported AUC. This table gives what they achieve on their own, under the same repeated stratified 5-fold pipeline. In the Miltiadous contrasts demographics alone reach an AUC near 0.64, so a portion of the roughly 0.85 AUC reported for every representation on those contrasts is demographic rather than neural, and the comparison between representations, which is what this paper is about, is unaffected because every representation carries the same demographic information. In the two Parkinson's cohorts demographics alone fall below chance. MOMENT and Chronos were originally scored without these predictors, because their exported matrices did not carry the demographic columns; they are now scored with them, so all nine representations use identical inputs.

| Cohort | Contrast | n | Age + sex only AUC |
|---|---|---|---|
| Miltiadous | AD vs HC | 34 v 29 | 0.639 |
| Miltiadous | FTD vs HC | 22 v 29 | 0.631 |
| Rockhill | PD vs HC | 15 v 16 | 0.246 |
| Cavanagh | PD vs HC | 25 v 25 | 0.326 |

### Supplementary Figures

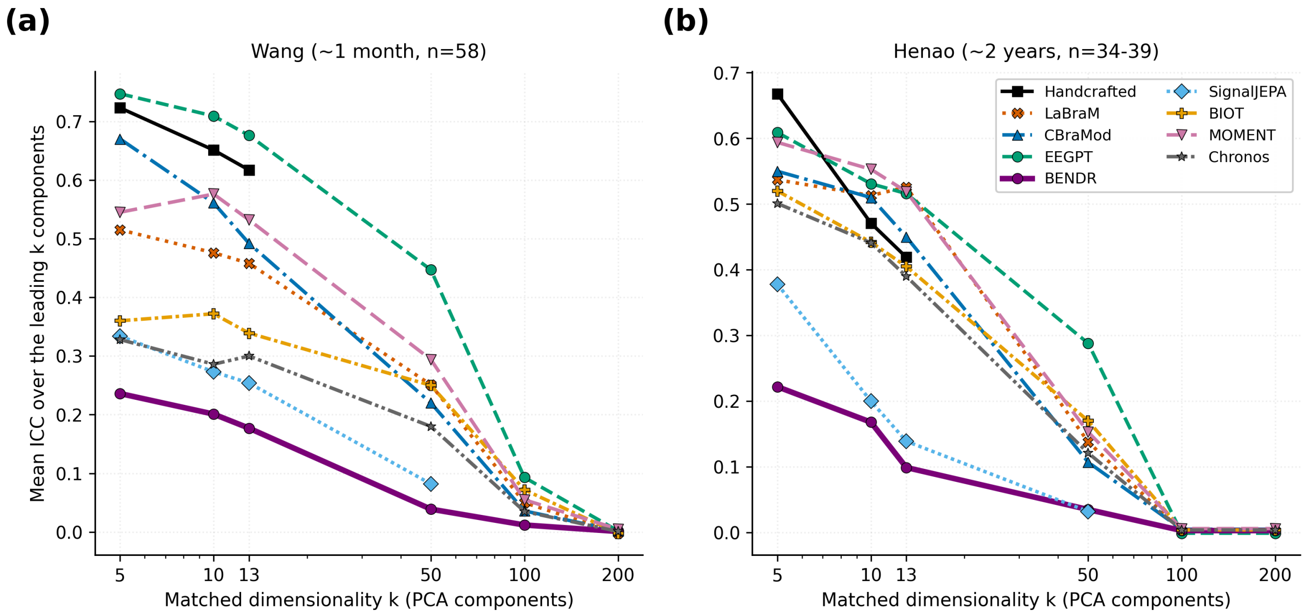


**Supplementary Figure S1. Matched-dimensionality reliability across principal components.**
Mean ICC over the leading k principal components for each representation, including MOMENT and Chronos, in the Wang cohort (a) and the Henao cohort (b). Holding dimensionality constant across representations removes the concern that the reliability ordering in Table 1 of the main text is an artifact of embedding size, which ranges

from 13 to 2048 across the nine representations. The reliable cluster remains above LaBraM, and BENDR remains lowest, across informative variance ranges. Underlying values are in **Supplementary Table S8**.

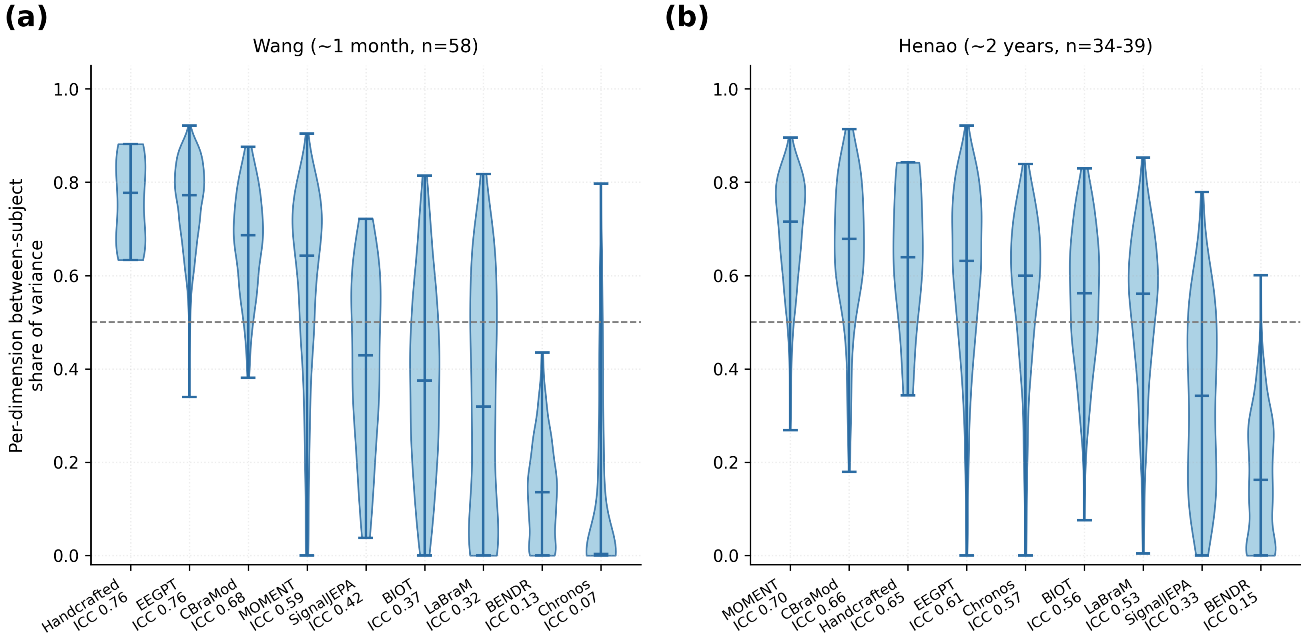


**Supplementary Figure S2. Between-subject share of variance, by representation and cohort.**
Violin plots show the distribution across embedding dimensions of the proportion of total variance attributable to between-subject differences, for all nine representations in the Wang cohort (a) and the Henao cohort (b). Horizontal bars mark the median and the range. Representations are ordered by mean ICC within each panel, and each is annotated with that value. The dashed line marks an equal split between the between-subject component and all other variance. Reliable representations concentrate variance in the between-subject component, whereas unreliable representations place a larger share into session and residual variance. This is the variance-level counterpart of the ICC ordering in Section 3.1, and it underlies the diagnostic used there to separate genuine reliability from variance compression under added noise.

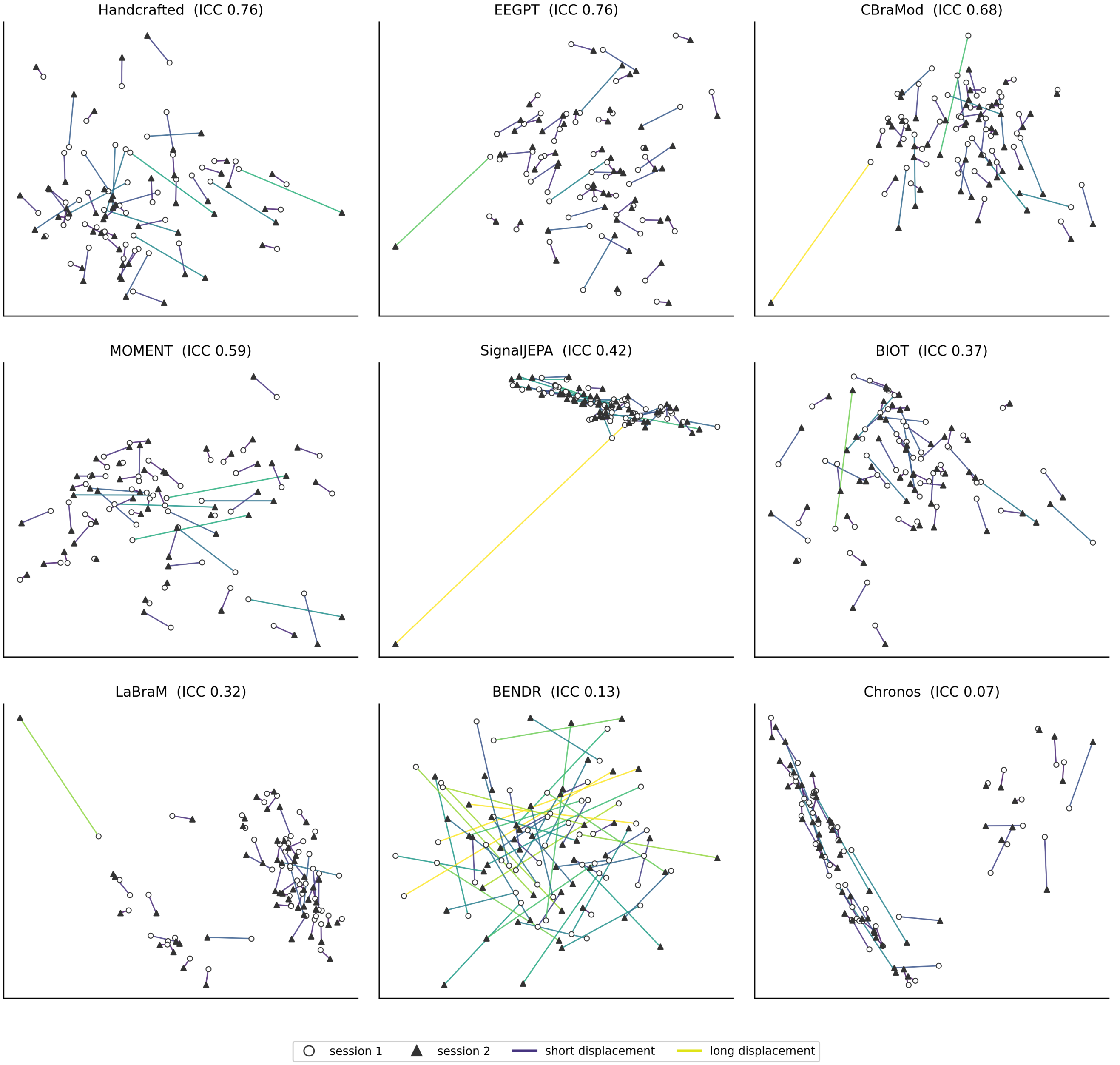


**Supplementary Figure S3. Session-to-session embedding drift trajectories, Wang cohort.**
Each line connects one subject's first and second session in a two-dimensional principal-component projection of that representation's embedding space, with open circles marking session 1 and filled triangles session 2. Panels are ordered by mean ICC, given in each title. Line colour encodes the size of a subject's session-to-session displacement relative to the spread of the cloud, so darker lines indicate a subject whose sessions stayed close together. Reliable representations produce short displacements relative to between-subject spacing, so a subject's sessions remain clustered, whereas unreliable representations produce displacements comparable to the distance between different subjects, which is most visible for BENDR. Axes are unlabelled because principal-component coordinates are arbitrary and differ between panels; only the relative geometry within a panel is meaningful. This is the geometric intuition behind the within-to-between distance ratio reported in Table 1 of the main text.

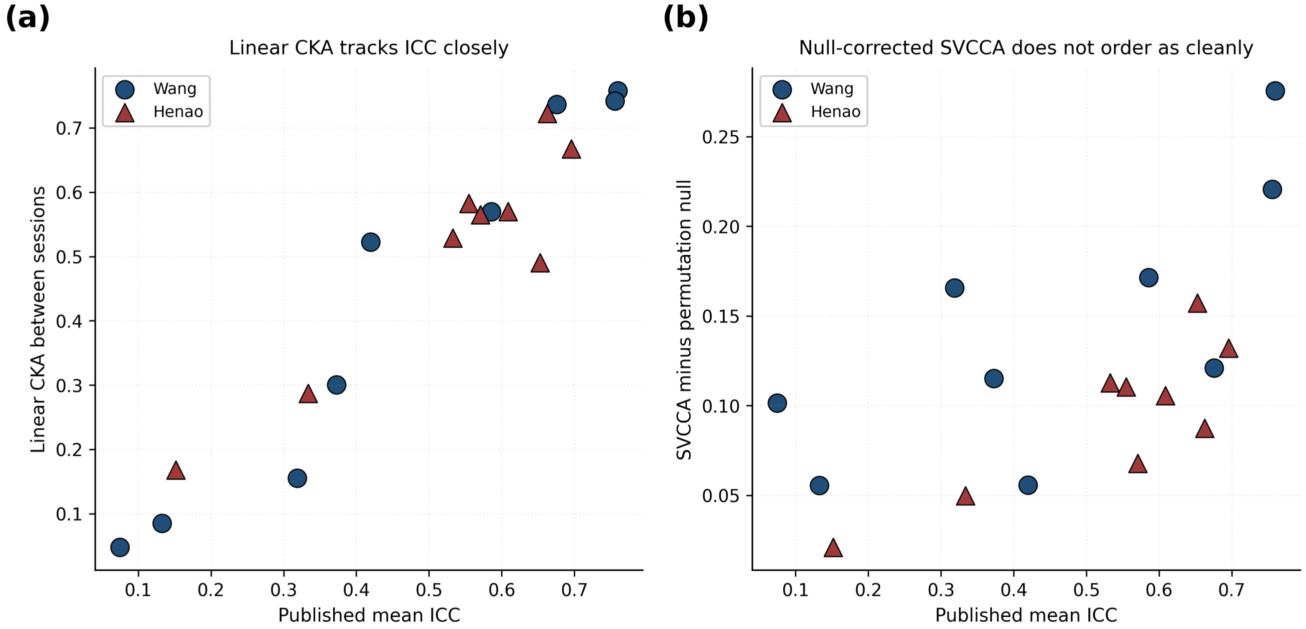


**Supplementary Figure S4. Representational similarity between sessions compared against reliability.** (a) Linear CKA between the same subject's sessions, plotted against that representation's mean ICC. (b) SVCCA similarity with its permutation null subtracted, plotted against the same quantity. Circles are the Wang cohort and triangles the Henao cohort. Linear CKA tracks ICC closely and monotonically across both cohorts, which is the independent confirmation referred to at the start of Section 3.2: a similarity measure built on entirely different assumptions recovers the same ordering. The null-corrected SVCCA values are positive for every representation but do not order them as cleanly, so we rely on the CKA comparison rather than the SVCCA one for that argument.

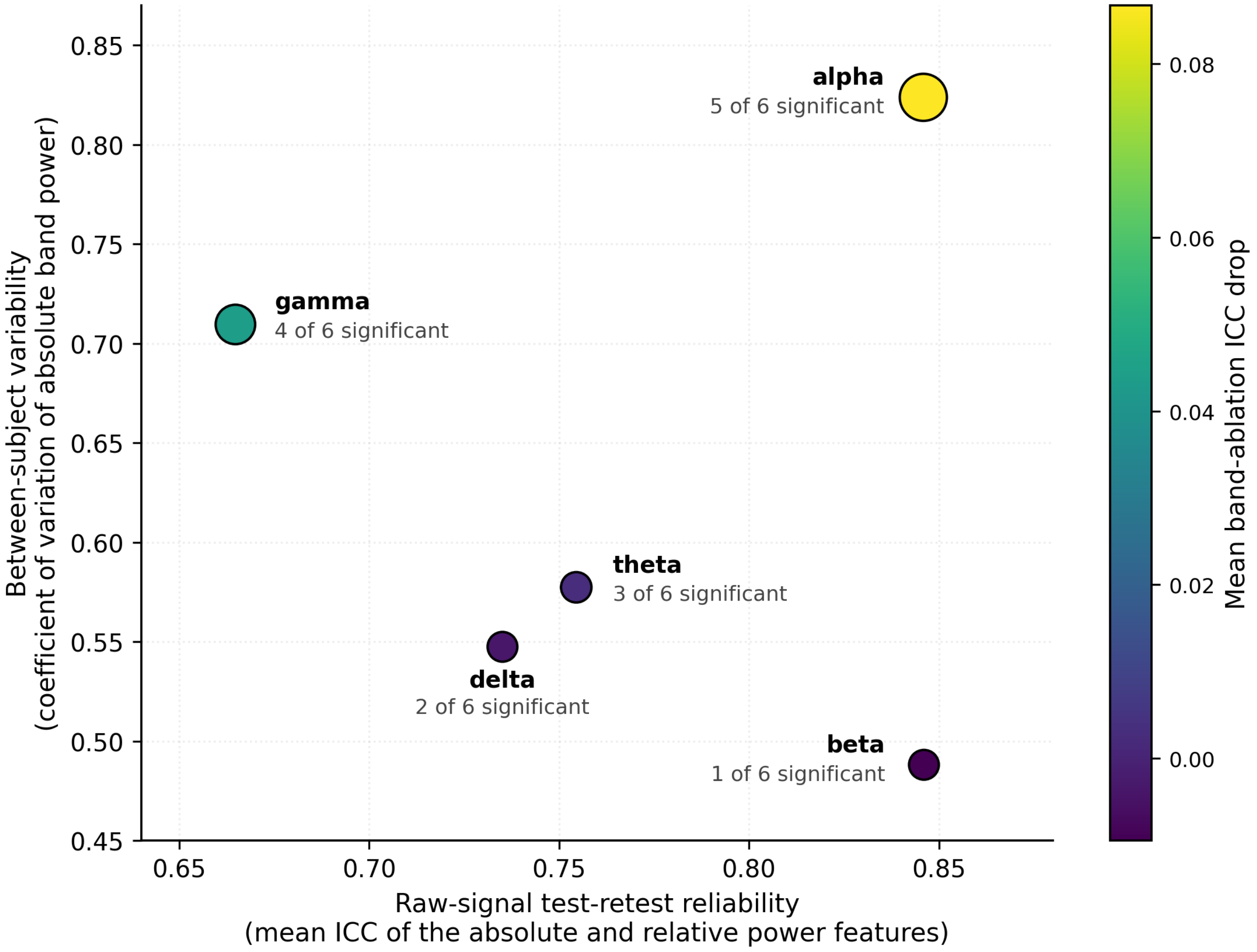


**Supplementary Figure S5. Band reliability, between-subject variability, and ablation importance compared.** Each point is one frequency band, positioned by its raw-signal test-retest reliability (x) and its between-subject coefficient of variation (y), with colour encoding the signed mean band-ablation ICC drop across all six perturbation-tested representations, EEGPT, CBraMod, LaBraM, BIOT, BENDR and Chronos, pooled irrespective of whether a given representation's own effects reached significance, and marker size emphasising positive mean drops, so that the two bands with negative means, beta and delta, are drawn at the same minimum size. The count in each label is the number of those six representations showing a significant effect for that band. Alpha occupies the upper right, combining high reliability with the highest between-subject variability, which is the pattern described in Section 3.2 of the main text. Beta sits at comparable reliability but the lowest variability and has negligible ablation importance. Gamma is the second-largest ablation driver despite the lowest raw-signal reliability of any band, and unlike alpha it is nearly irrelevant to disease discrimination; the muscle-artifact hypothesis offered for this in Section 3.2 rests only on indirect proxies and is not established here.
